# The role of *polycystic kidney disease-like* homologs in planarian nervous system regeneration and function

**DOI:** 10.1101/2024.07.17.603829

**Authors:** Kelly G. Ross, Sarai Alvarez Zepeda, Mohammad A. Auwal, Audrey K. Garces, Sydney Roman, Ricardo M. Zayas

**Author notes:** **Correspondence:** Ricardo M. Zayas.

## Abstract

Planarians are an excellent model for investigating molecular mechanisms necessary for regenerating a functional nervous system. Numerous studies have led to the generation of extensive genomic resources, especially whole-animal single-cell RNA-seq resources. These have facilitated *in silico* predictions of neuronal subtypes, many of which have been anatomically mapped by in situ hybridization. However, our knowledge of the function of dozens of neuronal subtypes remains poorly understood. Previous investigations identified that *polycystic kidney disease (pkd)-like* genes in planarians are strongly expressed in sensory neurons and have roles in mechanosensation. Here, we examine the expression and function of all the *pkd* genes found in the *Schmidtea mediterranea* genome and map their expression in the asexual and hermaphroditic strains. Using custom behavioral assays, we test the function of *pkd* genes in response to mechanical stimulation and in food detection. Our work provides insight into the physiological function of sensory neuron populations and protocols for creating inexpensive automated setups for acquiring and analyzing mechanosensory stimulation in planarians.

## Introduction

Freshwater planarians are an excellent organism to study stem cell-based regeneration due to their large population of stem cells and prodigious ability to regenerate all cell types, including neurons (Reddien, 2018; Ivankovic et al., 2019). The planarian nervous system is anatomically simple, but studies have demonstrated the neuronal population is highly heterogeneous, with many discrete cell types that are spatially restricted and give rise to form and function (Ross et al., 2017). From early estimates of dozens of specialized neuronal cell types, we do not know how many neuronal cell types or the exact functions of those cells in planarians (Fincher et al., 2018; Plass et al., 2018; King, H. O. et al., 2024). To fully comprehend how planarians regenerate neurons and restore function, generating a molecular neuroanatomical map and introducing tools to examine neuronal function will be critical. Efforts to create single-cell gene expression atlases have generated planarian cell-type profiles (Fincher et al., 2018; Plass et al., 2018; King, H. O. et al., 2024). Those studies have created two broad categories of neurons, ciliated and non-ciliated neurons, comprising the planarian nervous system (Fincher et al., 2018). Among the most prominent genes defining neural cells are *polycystic kidney disease-like* genes that define discrete ciliated neuronal populations, some of which have been implicated in mechanosensory and chemosensory functions (Fincher et al., 2018; Ross et al., 2018; Arnold et al., 2021).

PKD proteins are among one of the most ancient classes of sensory receptors. They are a member of the transient receptor potential (TRP) family of ion channels, which can be grouped into subfamilies based on the type of stimulus they are receptive to. The infamous names of these genes come from their roles in autosomal dominant polycystic kidney disease (ADPKD) caused by mutations of the PKD1 or PKD2 genes that encode Polycystin-1 (PC1) and Polycystin-2 (PC2) proteins (Harris and Torres, 2009; Esarte Palomero et al., 2023). The rest of the PKD1 and −2 gene families consist of PKD1-LIKE and PKD2-LIKE proteins which are identified and grouped based on their structural and sequence similarities to the founding PC1 and −2 proteins (Esarte Palomero et al., 2023). PKD1 and PKD2 family members are integral membrane proteins that can exist as heterotetrameric receptor-like/ion channel complexes that transduce signals by conducting Ca^2+^ currents or potentially acting as a G-protein coupled receptor (Maser et al., 2022; Esarte Palomero et al., 2023). Multiple combinations of PKD1 family and PKD2 family members have been found in these heterotetramers creating a diverse array of complexes (Esarte Palomero et al 2023). This variety of complexes along with subcellular distribution - these proteins are found in multiple cellular locations, like the ER and primary cilia - implicate these proteins in diverse cellular processes in sensory neuron function, like mechanosensation and chemosensation, signal transduction and gene expression, gustatory sensing, and the sperm acrosome reaction (Esarte Palomero et al., 2023). Additionally, the evolutionary history and function of PKD genes is understudied. Homologs of PC1 proteins have not been found in invertebrate organisms and might represent vertebrate-specific gene duplication and evolution events. Thus, understanding the ancestral roles of PKD genes will benefit from studies in diverse organisms.

Most animals possess homologs of Pkd1- and Pkd2-like genes. Evidence from invertebrate model organisms, including acoels, planarians, *C. elegans*, *Hydra*, and *Nematostella vectensis*, indicate Pkd-like genes are expressed in nervous system cells (Barr et al., 2001; O’Hagan et al., 2014; McLaughlin, 2017; Fincher et al., 2018; Sebe-Pedros et al., 2018; Hulett et al., 2024; Sakagami et al., 2024). TRP channels have been noted to have an evolutionarily conserved role in mechanosensation and are present in a variety of organisms and clades (Liu and Montell, 2015). Across taxa, there is extensive variability in the roles of PKD channels. For example, Polycystin-2 can function without Polycystin-1 or be regulated by other receptors. In fission yeast, a Pkd2 channel functions alone to sense the change in pressure during cytokinesis at the cleavage furrow and is required for proper cell division (Morris et al., 2019). In addition, *Drosophila* Pkd2 participates in phagocytosis of apoptotic cells (Van Goethem et al., 2012). In several sexually reproducing organisms, *pkd2* is expressed in sexual organs. In *C. elegans*, the Pkd-1 homolog *lov-1* and *pkd-2* are responsible for proper mating function in males (Barr et al., 2001; O’Hagan et al., 2014). In mice, there may be a role for *Pkd1L-3* and *Pkd2L-1* in the taste buds for sour taste detection (Ishimaru et al., 2006; Horio et al., 2011).

In planarians, Pkd genes are not expressed in the excretory organs (Thi-Kim Vu et al., 2015). Evidence from several studies indicates that Pkd1- and Pkd2-like genes, as well as other TRP genes, are expressed in sensory neurons (Inoue et al., 2014; Fincher et al., 2018; Ross et al., 2018; Arnold et al., 2021). Because of their striking expression patterns, abundant transcript levels, and essential roles in diverse cell physiology processes, we sought to further characterize *pkd* homologs and examine unexplored family members. We performed comprehensive expression experiments in *S. mediterranea pkd* family genes and showed that they are expressed in neurons in asexual planarians. Moreover, expression analyses in sexually reproducing planarians revealed expression in the reproductive structures, suggesting *pkd* genes might also play roles in reproduction. We assessed *pkd* gene function using custom assays to characterize the physiological roles of *pkd^+^* sensory neurons in planarians. Furthermore, we describe simple setups for performing quantitative behavioral assays for mechanosensation and chemosensation. Our findings contribute to our understanding of planarian regeneration and neurobiology and will be useful for comparative studies of *pkd-like* gene evolution.

## Results

### Planarian PKD Protein Domain Structure

To determine if additional *pkd* genes were present in *S. mediterranea*, we extracted the highly conserved PKD domains from *pkd1L-2* and *pkd2L-1* as representatives of the *pkd1* and *pkd2* gene families, respectively, and ran BLAST searches against the planarian transcriptome (Rozanski et al., 2019). The searches revealed all nine known *pkd* transcripts, including six genes that we previously examined in (Ross et al., 2018) (Table S1), which is also consistent with the *pkd* homologs identified in (Thi-Kim Vu et al., 2015). We noted that two of these genes were categorized as alternative forms of the same transcript in the dd_Smed_v6 transcriptome (*pkd2L-*2: dd_v6_17348_0_3 and *pkd2L-3*: dd_v6_17348_0_2). However, these sequences do not align with each other and each sequence aligns with separate contigs in the *S. mediterranea* genome (Grohme et al., 2018). These transcripts might represent two unique genes. Bayesian phylogenetic analyses support that the planarian *pkd* genes are clear homologs of PKD1-like (Fig. S1) and PKD2 and PKD2-like proteins (Fig. S2).

We used PFAM to compare the domains in planarian PKD1 and PKD2 predicted proteins to the four PKD1 family proteins and three PKD2 (TRPP) family proteins found in humans. PKD2 family proteins are much smaller than PKD1 family proteins in general and *S. mediterranea* predicted proteins follow this trend similarly to the human proteins (Fig. 1A). Human PC1 is a larger protein than all other PKD-like proteins found in humans and planarians alike; PC1 is 4303 AA in length, more than 1.5X the size of the next largest human PKD family gene (PKD1L1), which is 2849 AA long (Fig 1A, Accession numbers in Methods). PC1 also contains several N-terminal extracellular domains located on the N-terminal side of the extracellular Receptor for Egg Jelly (REJ) domain that are not found in other PKD family proteins (Fig. 1A). Following these extracellular domains are eleven transmembrane domains with an intracellular polycystin-1, lipoxygenase, and α-toxin (PLAT) domain that follows the first transmembrane domain. This domain was found in all human and planarian PKD sequences. A portion of the transmembrane sequence is classified as the highly conserved Polycystin Domain based on similarity to the PC1 sequence; this region is found in all PKD1 and −2 proteins in both humans and planarians. Finally, regions within both the PKD1 and −2 family proteins contain a highly conserved sequence known as the PKD cation channel (Fig. 1). Interestingly, although PKD1 proteins were historically known to function as receptors that cluster with one or more PKD2 genes that act as the cation channel effector of this complex, recent research has shown that the last six transmembrane domains of PKD1 proteins are functioning as part of the cation channels (Esarte Palomero et al., 2023). There is evidence that PKD1 proteins form complexes with multiple PKD proteins (Esarte Palomero et al., 2023); given the conservation of domains seen between human and planarian PKD genes, it is possible that planarian PKD1 proteins form complexes with multiple PKD2 proteins as well.

**Figure 1.**
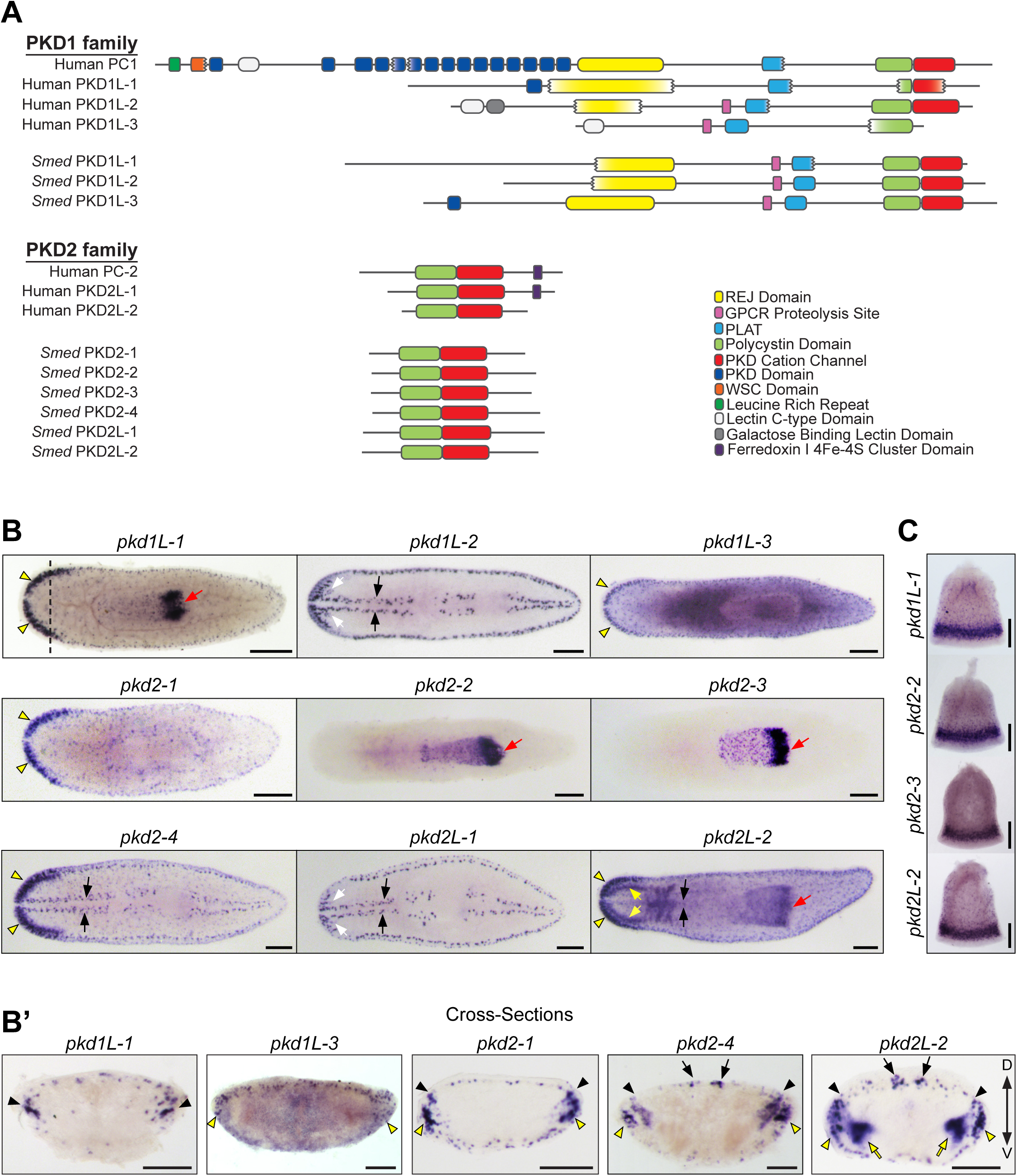
Predicted domain structures of human and planarians and whole-mount in situ hybridizations in asexual *Schmidtea mediterranea*. The protein domains of PKD1 and PKD2, human and planarian proteins, illustrate the conserved presence of key structural and functional domains in these proteins. REJ, Receptor for Egg Jelly; GPCR, G-Protein Coupled Receptor; PLAT, Polycystin-1, Lipoxygenase, and alpha toxin; WSC, cell wall integrity and stress response component. The exception is the PKD1 (Polycystin-1) protein, which has been only found in chordates and possesses a complex N-terminal extracellular domain. (B) *pkd* gene WISH in asexual planarians reveals a variety of expression patterns, including expression in dispersed subepidermal cells, in the dorsal ciliated stripe and peripheral stripes (black arrows), in the pharynx (red arrows), photoreceptors (white arrows), in the auricles and brain branches (yellow arrowheads), and in the brain (yellow arrows). Scale bars, 200 µm. (B’) Cross-sections of WISH from (B) of brain branch- and auricle-expressed *pkd* genes highlight expression in either the brain branches (black arrowheads), the auricle region (yellow arrowheads), and the brain (yellow arrows). Expression in the dorsal ciliated stripe is highlighted with black arrows. Scale bars, 100 µm. (C) WISH on isolated pharynges for genes with pharyngeal expression shows the expression of *pkd* transcripts in discrete puncta throughout the pharynx and abundant expression in the pharyngeal nerve net at the distal end (bottom of each image). Scale bars, 50 µm. n ≥ 8 worms tested, with all worms displaying similar expression patterns for all genes.

### Planarian *pkd-like* genes are expressed in neurons

To examine the expression of *pkd* genes in *S. mediterranea*, especially for the three untested genes, we performed whole-mount in situ hybridization (WISH) on intact asexual planarians. We observed labeling in multiple sensory-rich regions for all nine *pkd* genes (Fig. 1B). We also performed transverse cross-sections through the head regions of planarians to improve the spatial resolution of *pkd1L-1, pkd1L-3, pkd2-1, pkd2-4,* and *pkd2L-2*, which were expressed in the auricles and brain branches (Fig1B’). Additionally, we performed WISH on chemically amputated pharynges to facilitate visualization of *pkd* labeling in pharyngeal tissues (Fig. 1C). Taken together, these experiments show that planarian *pkd* genes are strictly expressed in seven sensory cell-type rich structures and patterns (summarized in Table 1). Our results are consistent with *pkd* gene in situ experiments in other studies. Following on our previous work (Ross et al., 2018), we observed the expression of *pkd1L-*2, *pkd2-4, pkd2L-1,* and *pkd2L-2* in the rheosensory organ, which is comprised of the dorsal ciliated stripe as well as the ventral and dorsal peripheral stripes (highlighted with arrows, Fig. 1B and 1B’). This population of cells represents submuscular neurons that are involved in sensing mechanical stimulation, such as water flow and vibration (Ross et al., 2018). *pkd1L-2* and *pkd2L-1* have the most discrete labeling patterns of all planarian *pkd* genes; they are expressed in the rheosensory organ and in a small population of photoreceptor neurons (Ross et al., 2018).

**Table 1.** Summary of *Schmidtea mediterranea* PKD-like gene expression patterns, regulation by *soxB1-2*, and RNAi phenotypes.

| Gene Name | Dresden Transcript ID | Expression Pattern<br>(Loss of expression observed after *soxB1-2* RNAi: <u>Yes</u>/<u>No</u>/<u>Partial</u>) |  |  |  |  |  |  | Observed Behavioral Phenotype |
| --- | --- | --- | --- | --- | --- | --- | --- | --- | --- |
|  |  | BB | Aur | Brain | RO | DSC | PR | Ph |  |
| **Pkd1 Family Members** |  |  |  |  |  |  |  |  |  |
| *pkd1L-1* | dd_Smed_v6_15525_0_1 | ✓<br>(Y) |  |  |  | ✓<br>(Y) |  | ✓<br>(N) | Reduced chemosensory behaviors |
| *pkd1L-2* | dd_Smed_v6_13975_0_1 |  |  |  | ✓<br>(Y) |  | ✓<br>(N) |  | Reduced mechanosensory behaviors |
| *pkd1L-3* | dd_Smed_v6_10962_0_1 |  | ✓<br>(Y) |  |  | ✓<br>(Y) |  |  | None |
| **Pkd2 Family Members** |  |  |  |  |  |  |  |  |  |
| *pkd2-1* | dd_Smed_v6_12955_0_1 | ✓<br>(Y) | ✓<br>(Y) |  |  | ✓<br>(Y) |  |  | None |
| *pkd2-2* | dd_Smed_v6_17348_0_3 |  |  |  |  |  |  | ✓<br>(N) | None |
| *pkd2-3* | dd_Smed_v6_17348_0_2 |  |  |  |  |  |  | ✓<br>(N) | None |
| *pkd2-4\** | dd_Smed_v6_13327_0_1 | ✓<br>(P) | ✓<br>(Y) |  | ✓<br>(Y) | ✓<br>(Y) |  |  | D26-D40, 8-feed RNAi:<br>Reduced chemosensory behaviors; Reduced mechanosensory behaviors<br>Extended RNAi, D55, 8- and 14-feed RNAi:<br>slow inching, jerky movements observed in Ross et al., 2018. (18/18 worms) |
| *pkd2L-1* | dd_Smed_v6_15626_0_1 |  |  |  | ✓<br>(Y) |  | ✓<br>(**) |  | Reduced mechanosensory behaviors |
| *pkd2L-2* | dd_Smed_v6_9977_0_3 | ✓<br>(P) | ✓<br>(Y) | ✓<br>(N) |  | ✓<br>(Y) |  | ✓<br>(N) | None |

Abbreviations: BB, Brain Branches; Aur, Auricles; RO, rheosensory organ (both the dorsal ciliated stripe and dorsal and ventral peripheral ciliated stripes); DSC, Dispersed Subepidermal Cells; PR, photoreceptors; Ph, pharynx; D, day; DPA, days post-amputation. \*The severity of movement defects precluded quantification of behaviors for the extended RNAi. Videos of control and *pkd2-4(RNAi)* D28 (normal movement) and D55 (abnormal movements) shown in Videos S1-S4. \*\*Not scored.

Five *pkd* genes, *pkd1L-1, pkd1L-3, pkd2-1, pkd2-4,* and *pkd2L-2,* were expressed in the brain branches or auricles (Fig. 1B., yellow arrowheads). These structures have been implicated in planarian chemosensation (Koehler, 1932; Fraenkel, 1961; Farnesi and Tei, 1980; Roberts-Galbraith et al., 2016; Almazan et al., 2021). Most of these genes were expressed in both anatomical regions (auricles, black arrowheads, and brain branches, yellow arrowheads in Fig 1B’) apart from *pkd1L-1* and *pkd1L-3,* which were only expressed in the brain branches and auricles, respectively (Fig. 1B’). Additionally, *pkd2L-2* expression extended well into the ventromedial brain (yellow arrows in Fig 1B and 1B’). Interestingly, all *pkd* genes with expression in the auricle or the brain branches were also expressed in populations of dispersed subepidermal cells. Expression of *pkd1L-3* was previously attributed to both neurons and epidermal cells in Ross et al. (2018) due to detectable transcript expression in a single *soxB1-2^+^*epidermal cell identified in (Wurtzel et al., 2015). However, newer data does not strongly indicate *pkd1L-3* expression in the epidermis (Fincher et al., 2018), which is consistent with our observations. Four *pkd* genes, *pkd1L-1, pkd2-2, pkd2-3,* and *pkd2L-2,* were expressed in the pharynx (red arrows, Fig. 1B and Fig. 1C). Expression patterns for *pkd2-2* and *pkd2-3* looked nearly identical and were exclusively detected in the pharynx. In contrast, *pkd1L-1* and *pkd2-4* had broader expression domains. All *pkd* genes expressed in the pharynx had a similar pattern of expression throughout the pharynx with abundant expression at the distal end of the pharynx, in the pharyngeal nerve net. We did not observe *pkd* labeling in the protonephridia, consistent with reporting in Thi-Kim Vu et al. (2015).

We next sought to determine if the gene expression patterns correlated with cell-type assignments revealed by the single-cell RNA-seq atlas of whole-body planarian cells (Fincher et al., 2018). When we examined the enrichment of the *pkd* transcripts on scRNA-seq tSNE plots of *S. mediterranea* ciliated neurons, we observed a connection between *pkd^+^* WISH patterns and expression across the single cell clusters (Fig. 2). First, the genes expressed almost exclusively in the rheosensory organ (*pkd1L-2* and *pkd2L-1*) had clear enrichment in a small ciliated neural population (black circles, Fig. 2B). Genes *pkd2-4* and *pkd2L-2*, which are also expressed in the rheosensory organ, also showed high levels of enrichment in this population (black circles in Fig. 2E). The transcript for *pkd2-2* was not present in the Fincher et al. dataset (Fincher et al., 2018), likely due to a discrepancy of this distinct transcript as an isoform of *pkd2-3*. We found that *pkd2-3* is spread in a series of subclusters (blue circle on the right side of the tSNE plot, Fig. 2D). The other *pkd* genes with expression in the pharynx (*pkd1L-1* and *pkd2L-2*) were similarly enriched throughout clusters in this subset of the plot (blue circles, Fig. 2E). *pkd1L-3* and *pkd2-1*, which were expressed within populations of the brain branches, auricles, and dispersed subepidermal cells, displayed enrichment in a subset of clusters likely representing different neuronal cell types (red circle, Fig. 2C); *pkd1L-1, pkd2-4,* and *pkd2L-2* also showed expression in these regions by WISH and the corresponding scRNA-seq populations amongst others (red circle, Fig. 2E). In our previous work, we showed that *pkd1L-2^+^* cells expressed pan-neural markers and the cilia gene, *rootletin* (Ross et al., 2018). Consistent with previous observations (Fincher et al., 2018), we detected co-enrichment of *rootletin* and pan-neuronal genes *synapsin* and *synaptotagmin* in the *pkd^+^* scRNA-seq clusters (Fig. 2F). We tested these observations by dFISH, which showed *pkd2L-2* is co-expressed with cilia and pan-neural markers (Fig. 2G).

**Figure 2.**
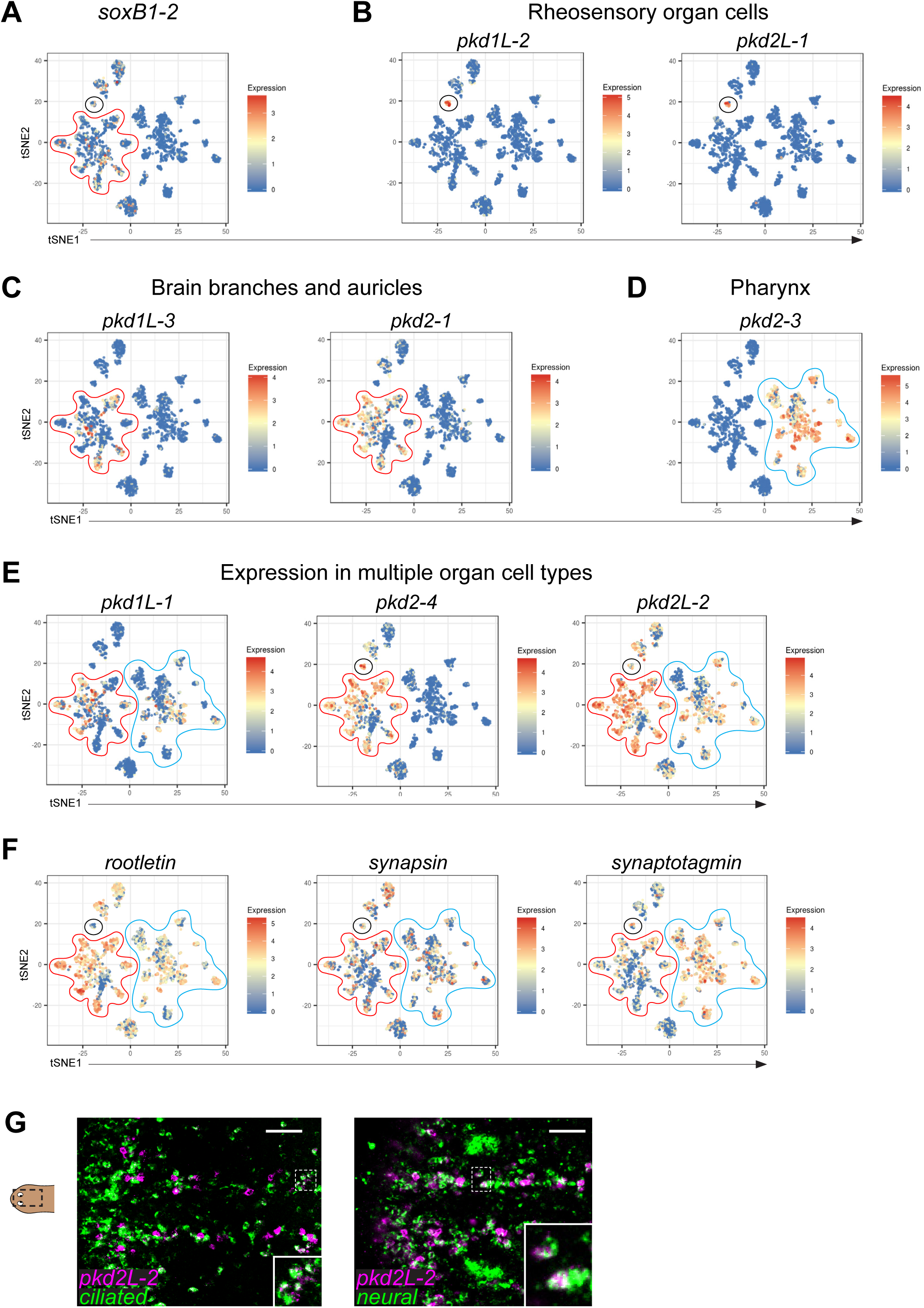
**Planarian scRNA-seq tSNE plots demonstrate that *pkd^+^* cells are ciliated neuronal cells and that a subpopulation of *pkd^+^* cells co-express *soxB1-2.*** (A-F) tSNE plots for all *pkd* genes were downloaded from https://digiworm.wi.mit.edu/ (Fincher et al., 2018). Each dot represents a single cell, with the relative expression of the gene represented as the color of the dot, as defined by the heatmap on the right side of each tSNE plot. (A) *soxB1-2* is expressed in clusters that contain the *pkd+* mechanosensory (rheosensory) population (black circle) and the *pkd+* chemosensory populations (clusters within the red outline). (B-F) tSNE plots for all the *pkd* genes except for *pkd2-2*, which is not represented in the Fincher et al. (2018) dataset. Broadly, *pkd* genes are expressed in one or more of three groupings of clusters which correspond to expression patterns seen in the whole animal (see Figure 2): the rheosensory organ cluster (black circle), the brain branch and auricle cluster (red outline), and the pharynx cluster (blue outline). (B) *pkd1L-2* and *pkd2L-1* are expressed in the rheosensory organ cluster (black circle). (C) *pkd1L-3* and *pkd2-1* expression is visualized in the brain branch and auricle clusters (red outline). (D) *pkd2-3* is expressed in the pharynx clusters (blue outline). (E) *pkd1L-1, pkd2-4,* and *pkd2L-2* are expressed in multiple clusters, as highlighted by outlining the pertinent clusters on the tSNE plots. (F) Co-expression of pan-neural (*synapsin* and *synaptotagmin*) and cilia (*rootletin*) marker genes in the *pkd^+^* clusters indicate that these cells are ciliated neurons. (G) to further confirm that co-expression of *synapsin*, *synaptotagmin*, and *rootletin* in *pkd^+^* clusters in the single cell data correlates with gene expression in the animal, the co-expression of these markers (green; *ciliated* = *rootletin, neural* = *synapsin/synaptotagmin* mix) with *pkd2L-2* (magenta) is shown. Dashed boxes represent the area of the inset in the images. Scale bars, 50 µm, n ≥ 3 worms tested with all samples displaying similar expression patterns.

Because PKD proteins have been implicated in mating behaviors and reproduction in other organisms (Kierszenbaum, 2004; Esarte Palomero et al., 2023), we were intrigued if *pkd* genes could be detected near or in the reproductive structures of the *S. mediterranea* hermaphroditic strain (Issigonis and Newmark, 2019). Thus, we performed WISH to *pkd* genes in sexual planarians and found that in addition to the striking expression patterns observed in the asexual worms, *pkd1L-1, pkd1L-3, pkd2-1, pkd2-2, pkd2-4, pkd2L-1, and pkd2L-2* were expressed in the copulatory apparatus (Fig. S3). Furthermore, *pkd1L-2* was expressed in the copulatory apparatus, oviduct, and adjacent to the glands. *pkd2-3* was expressed in the copulatory apparatus and the testes; we also noted that *pkd2-1* and *pkd2-4* showed expression in the gonopore area. We do not know if the genes are expressed in reproductive cells or if neurons are adjacent to reproductive structures, but the robust expression suggests it will be interesting to investigate how *pkd* function impacts reproduction in *S. mediterranea* hermaphrodites. Taken together, the expression patterns of planarian *pkd* genes demonstrate enrichment of expression in putative and previously characterized sensory neural populations.

### Analysis of *pkd* gene expression regulation by SoxB1-2

The *pkd^+^* sensory neuron clusters in the scRNA-seq plots suggest that these neuronal populations arise from divergent transcriptional regulatory networks. SoxB1-2 regulates six of the nine *pkd* genes (Ross et al., 2018); thus, in addition to previously characterized planarian *pkd* genes, we examined if *pkd1L-1*, *pkd2-2* and *pkd2-3* were impacted by *soxB1-2* activity. First, we analyzed *soxB1-2* expression in the single cell atlas and found that *soxB1-2* was expressed in the *pkd^+^*cluster group that represented brain branches and auricle expression as well as the cluster that represented rheosensory organ expression (red and black circles in Fig. 2A). *soxB1-2* expression was not detected in the group of clusters that corresponded with expression in the pharynx (Fig. 2D and 2E, blue circles). The six *pkd* genes that were previously identified as having reduced expression after *soxB1-2* RNAi had expression in both the red and black circled clusters. Thus, we investigated the impact of SoxB1-2 on *pkd* gene expression by performing RNAi for *soxB1-2* followed by WISH for *pkd* transcripts in both intact and regenerating fragments (RNAi scheme summarized in Fig 3A; results summarized in Table 1).

**Figure 3.**
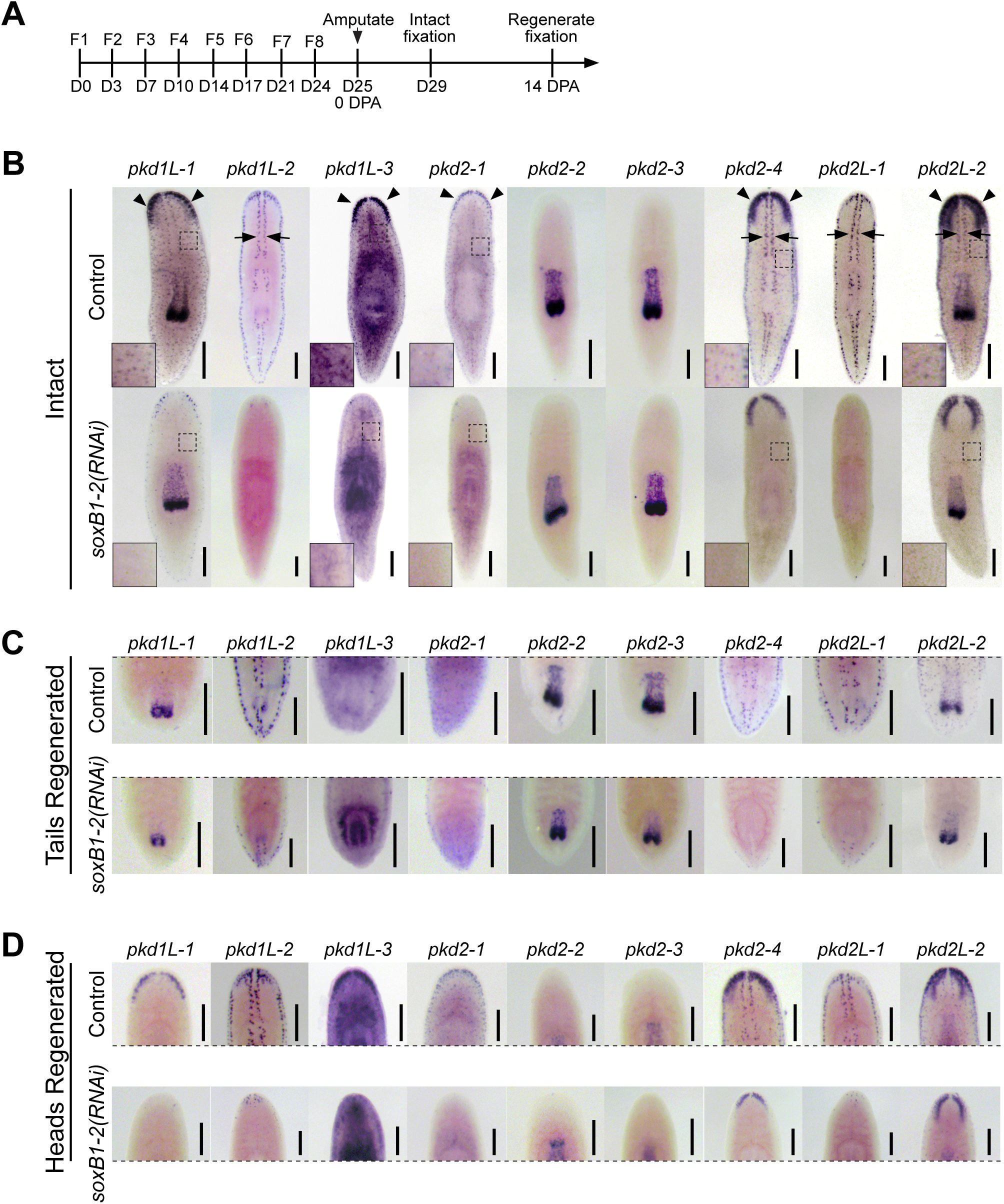
SoxB1-2 positively regulates *pkd* gene expression in sensory populations. (A) Timeline for assaying *pkd* gene expression by WISH following *soxB1-2* RNAi. (B-D) WISH for *pkd* genes on intact or 14 dpa *soxB1-*2(*RNAi*) regenerates that were amputated pre-pharyngeally. (B) Disrupting *soxB1-2* function in intact planarians led to reduced expression of *pkd* genes in the rheosensory organ (black arrows), the auricles, populations in the brain branches (black arrowheads), and in the dispersed subepidermal cells (inset images from dashed box areas). *pkd* expression appeared unaffected by *soxB1-2* RNAi in the pharynx and some brain branch populations as well as in the photoreceptors [see labeling in worms labeled with *pkd1L-1* (pharynx expression), *pkd1L-2* (photoreceptor expression), *pkd2-2* and *pkd2-3* (pharynx), *pkd2-4* (brain branch), and *pkd2L-2* (pharynx and brain branch)]. (C-D) *soxB1-*2*(RNAi)* head fragments that regenerated posterior tissues and new pharynx (C) or new heads (D) displayed loss of *pkd* labeling observed in intact planarians treated with dsRNA. Images cropped to display the regenerated tissues in each worm 14 dpa. Scale bars, 200 µm. n ≥ 3 worms tested, with all worms displaying similar expression patterns for all genes.

RNAi targeting *soxB1-2* resulted in a near to complete loss of *pkd* gene expression in the auricles (where *pkd1L-3, pkd2-1, pkd2-4, and pkd2L-2* are detected, black arrowheads in Fig. 3B), the rheosensory organ (arrows, Fig. 3B), and the dispersed subepidermal cells (insets in Fig. 3B). Loss of *soxB1-2* function resulted in the loss of *pkd1L-3* and *pkd2-1* expression in nearly all areas of the brain branches but resulted in the loss of *pkd2-4* and *pkd2L-2* expression only in a subset of cells in the brain branches. There was no apparent loss of expression in the ventromedial brain region in *pkd2L-2-*labeled worms or in the photoreceptors in *pkd1L-2-*labeled worms. Finally, *soxB1-2* RNAi did not affect *pkd* gene expression in the pharynx (where *pkd1L-1, pkd2-2, pkd2-3,* and *pkd2L-2* were detected). These results in intact worms were consistent in regenerating worms, except for *pkd1L-2* and *pkd2L-1* detection in a few anterior and posterior regenerated cells (Fig. 3C-D). Therefore, two of the *pkd* genes that were not previously identified as downstream of *soxB1-2* activity were, indeed, unaffected by *soxB1-2* RNAi; expression of *pkd2-2* and *pkd2-3* in the pharynx were comparable between controls and *soxB1-2(RNAi)* worms. This is consistent with the lack of overlapping expression gleaned from the scRNA-seq dataset (Fig. 2). Interestingly, *pkd1L-1* expression was downregulated following *soxB1-2* RNAi, but only in the brain branches and dispersed subepidermal cells and not the pharynx, which might explain why it was not detected as a differentially expressed gene in our published *soxB1-2* RNA-seq dataset (Ross et al., 2018).

### A subset of PKD genes is required for mechanosensation

Past studies, including classic literature, indicated that the dorsal ciliated stripe functions as a water-sensing organ (rheosensing), i.e., as a mechanosensing organ. At present, it is impractical to assess the neurophysiological properties of most neurons in planarians; therefore, we designed behavioral tests that could be used as readouts of impaired neurophysiological activity. Thus, we previously showed that disrupting *pkd1L-2* and *pkd2L-1* function using RNAi impairs rheosensation and vibration sensation (Ross et al., 2018). In addition, we described abnormal locomotion resulting from *pkd2-4* RNAi, including slow, inching, jerky movements that precluded inclusion in further behavioral assays. As a result, we sought to modify our RNAi scheme so that the onset of the *pkd2-4* RNAi phenotype requires longer to manifest, allowing us to perform behavioral analysis. Using this scheme (see Fig. S4A), we recorded animal movement defects by day 55 of the RNAi protocol; normal movements were recorded throughout behavioral assays (see reduced and extended feeding schemes and testing times in Fig. S4A-B and VideosS1-4 of normal movement for behavioral assays and extended feeding movement defects).

To determine if any additional *pkd* genes are required for mechanosensation, we employed the Ross et al. (Ross et al., 2018) vibration assay with modifications (see Methods and Fig. S4C-D). Control worms reliably contracted their bodies along the anteroposterior axis following a tapping stimulus; we measured responses by calculating the percent change in body length between pre-stimulus gliding length and post-stimulus length. We found that *pkd1L-2, pkd2L-1,* and *pkd2-4* RNAi displayed a significant decrease in response to the stimulus, while all other *pkd* genes had a similar contraction response as the controls (Fig. 4). We previously found that worms with a reduced ability to sense vibration in the water had a decreased ability to sense water flow applied across their dorsal side and, as with the vibration assay, lacked a normal contraction response (Ross et al., 2018). Therefore, we performed a rheosensory assay for the *pkd* genes showing a vibration defect (Fig. S4E). As an additional control, we also analyzed *pkd1L-3*, which is not expressed in the rheosensory organ. We found that the same three genes recapitulated a mechanosensory defect phenotype: *pkd1L-2, pkd2L-1,* and *pkd2-4* (Fig. S4F-G). Each of these transcripts was expressed in the dorsal ciliated stripe, implicating them as essential for the mechanosensory function of these neurons.

**Figure 4.**
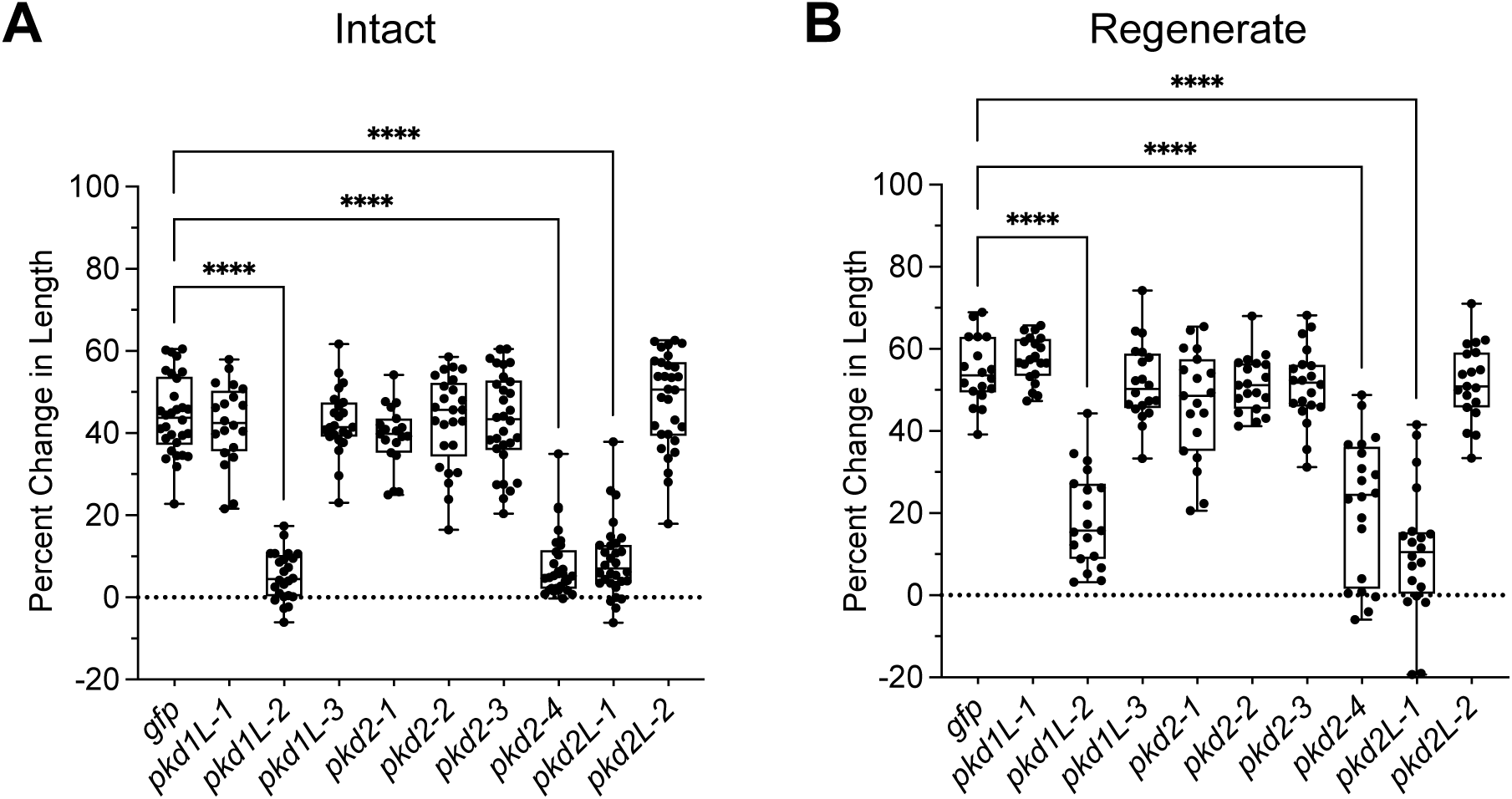
Behavioral analysis reveals *pkd* genes involved in detecting mechanosensory stimulation. (A-B) RNAi worms demonstrated significant reductions in their contraction behavior, represented as the percent change in length between pre-stimulus and post-stimulus worm length in vibration assays (see Figure S4 for the schematic of RNAi and Assay) in both intact (A) and regenerate (B) RNAi worms. *pkd1l-2*, *pkd2-4*, and *pkd2L-1* RNAi results in decreased mechanosensation. Each worm assayed is represented as a dot on the graph; the boxes represent data ranges from the 25^th^ to the 75^th^ percentile, with a bisecting line at the median and whiskers encompassing the full range of values for each experimental group. n ≥ 18 worms for each experimental group. ^∗∗∗∗^p < 0.0001; all other groups are non-significant (p > 0.01); One-way ANOVA with Dunnett’s multiple comparisons test.

### PKD genes required for chemosensation

Studies in vertebrate and invertebrate models implicate *pkd* gene function in chemosensory functions (LopezJimenez et al., 2006; Horio et al., 2011; O’Hagan et al., 2014). We observed the expression of five planarian *pkd* genes in the auricles and brain branches and the expression of four *pkd* genes in the pharynx (structures linked to chemosensory functions) (reviewed in Ross et al., 2017; Miyamoto et al., 2020). Thus, we investigated if planarian *pkd* gene function may be required for chemosensation. In our previous work, we noted reductions in the time *pkd2-1* and *pkd2L-1* RNAi worms spent in a ‘food zone’ of a circular dish, but without statistical confidence (Ross et al., 2018). To improve our analysis of chemosensory behaviors, we modified our assay (Fig. S5 and Methods). Using the modified assay, we found that of the intact worm groups, only *pkd2-4(RNAi)* worms showed a significant decrease in the time spent in the food zone. No groups showed a significant difference in the percent of worms that ate during the assay (Fig. 5A). However, in regenerates, both *pkd1L-1(RNAi)* and *pkd2-4(RNAi)* worms showed significant decreases in both the time in food zone and the percent of worms that ate during the assay (Fig. 5B). Both genes were expressed in the brain branches and *pkd2-4* was also expressed in the auricle area, indicating that cells within these structures are involved in chemosensing. While *pkd1L-1* is additionally expressed in the pharynx, other pharynx-expressed *pkd* genes had no significant feeding defects; we do not know at this time what role these *pkd* genes might be playing in mechanosensation or chemosensation in the pharynx. However, the overall reduced time in the food zone seen with loss of function of *pkd1L-1* and *pkd2-4* suggests the RNAi-treated planarians are unable to detect food in the water and hence fail to track the pellets in the first place, which attributes this phenotype to chemosensation. There are known roles for *pkd* genes in taste in other organisms (LopezJimenez et al., 2006; Horio et al., 2011), which might explain the abundant presence of some *pkd* genes in the tip of the pharynx; however, our experiments did not uncover defects that could be attributed to pharyngeal gene expression.

**Figure 5.**
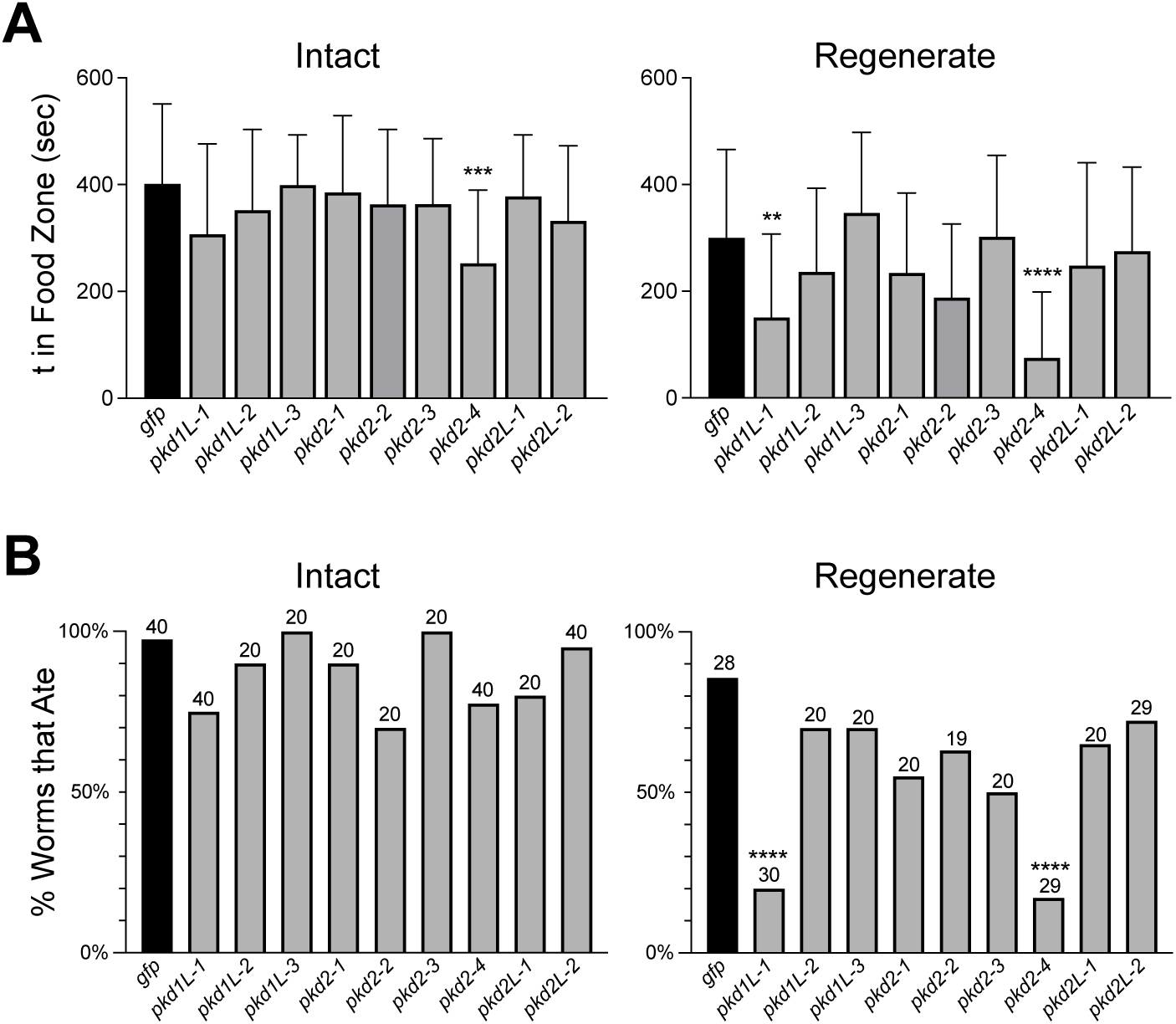
Feeding behavior analysis identifies candidate *pkd* genes implicated in chemosensation. (A-B) A chemosensation assay was performed on intact and regenerated worms at 14 dpa. (A) The average total time (t) intact and regenerate planarians spent in the ‘Food Zone’ (a 10 mm region surrounding liver pellets in the test field) during the 10-minute assay period demonstrated a significant reduction in time spent in the food zone for *pkd2-4* worms (intact) and both *pkd1L-1* and *pkd2-4* regenerate worms. Animals that did not enter the food zone during the assay period were assigned a value of 0. (B) The percent of the worms in each experimental group that ate in the 10-minute assay period was calculated for both intact worms and regenerates, demonstrating significant reductions in the number of eating *pkd1L-1*(*RNAi*) and *pkd2-4*(*RNAi*) regenerated worms. The total number of worms used for the assay is displayed above the bars and applies to the data for both assays (A). ^∗∗^p < 0.01, ^∗∗∗^p < 0.001, ^∗∗∗∗^p < 0.0001, all other values were determined as not significant (p > 0.01), One-way ANOVA with Dunnett’s multiple comparisons test.

### Co-expression of PKD1 and PKD2 family members in sensory neurons

PKD1 and PKD2 proteins are known to work cooperatively in cilia to facilitate sensory signal input, with PKD1 proteins acting as a receptor and PKD2 proteins acting as an ion channel (Esarte Palomero et al., 2023). We observed reduced chemosensing behavior with loss of *pkd1L-1* and *pkd2-4* expression (Fig. 5) and reduced mechanosensation when *pkd1L-2, pkd2L-1,* and *pkd2-4* were inhibited individually (Fig. 4). Because of the overlapping expression of *pkd1L-1* and *pkd2-4* in multiple scRNA-seq clusters (red circle, Fig. 2E) and expression of all three *pkd* genes that resulted in mechanosensory defects in the rheosensory organ scRNA-seq cluster (black circle, Fig. 2B and 2E), we reasoned that genes expressed in the same scRNA-seq clusters with behavioral phenotypes likely function in the same cells. Due to the known relationship between PKD1 and PKD2 proteins, we sought to determine if the pairs of *pkd1-like* and *pkd2* genes resulting in behavioral phenotypes were co-expressed using double-fluorescent in situ hybridizations (dFISH).

First, we performed dFISH on *pkd1L-1* and *pkd2-4* and observed co-expression in cells throughout the brain branch region in both ventromedial and dorsolateral cells (arrows and inset image in Fig 6A ventral and dorsal). *pkd2-4* labeling was much more abundant, which is in line with the increased number of cells that appear to have expression in both the WISH labeling (Fig. 1) and scRNA-seq tSNE plots (Fig. 2). Thus, it was not surprising that we were also able to see multiple cells in the head region that were only labeled with *pkd2-4* and not *pkd1L-1* (arrowheads in Fig. 6). Next, we examined co-labeling of the *pkd1 and −2* genes that resulted in mechanosensory phenotypes. *pkd1L-2* had very discrete labeling patterns by WISH and in the scRNA-seq plots, almost exclusively in the rheosensory organ (except for faint photoreceptor staining). However, we know of at least two distinct populations of cells in the rheosensory organ (one marked by *pkd1L-2* and the other by *hemicentin-1-like*) (Ross et al., 2018). So, we were uncertain whether these *pkd^+^* cells represented the same or distinct populations. We performed dFISH with these genes and found that these genes were, indeed, labeling the same discrete population of cells (white arrows and inset, Fig. 6). We also examined if *pkd2-4* was expressed in this cell population and found that *pkd1L-2* cells expressed *pkd2-4* (Fig. 6B, arrows); we again observed many additional cells labeled solely by *pkd2-4* (Fig. 6B, arrowheads). Thus, our data suggest that PKD1 and PKD2 genes function together in subsets of sensory cells.

**Figure 6:**
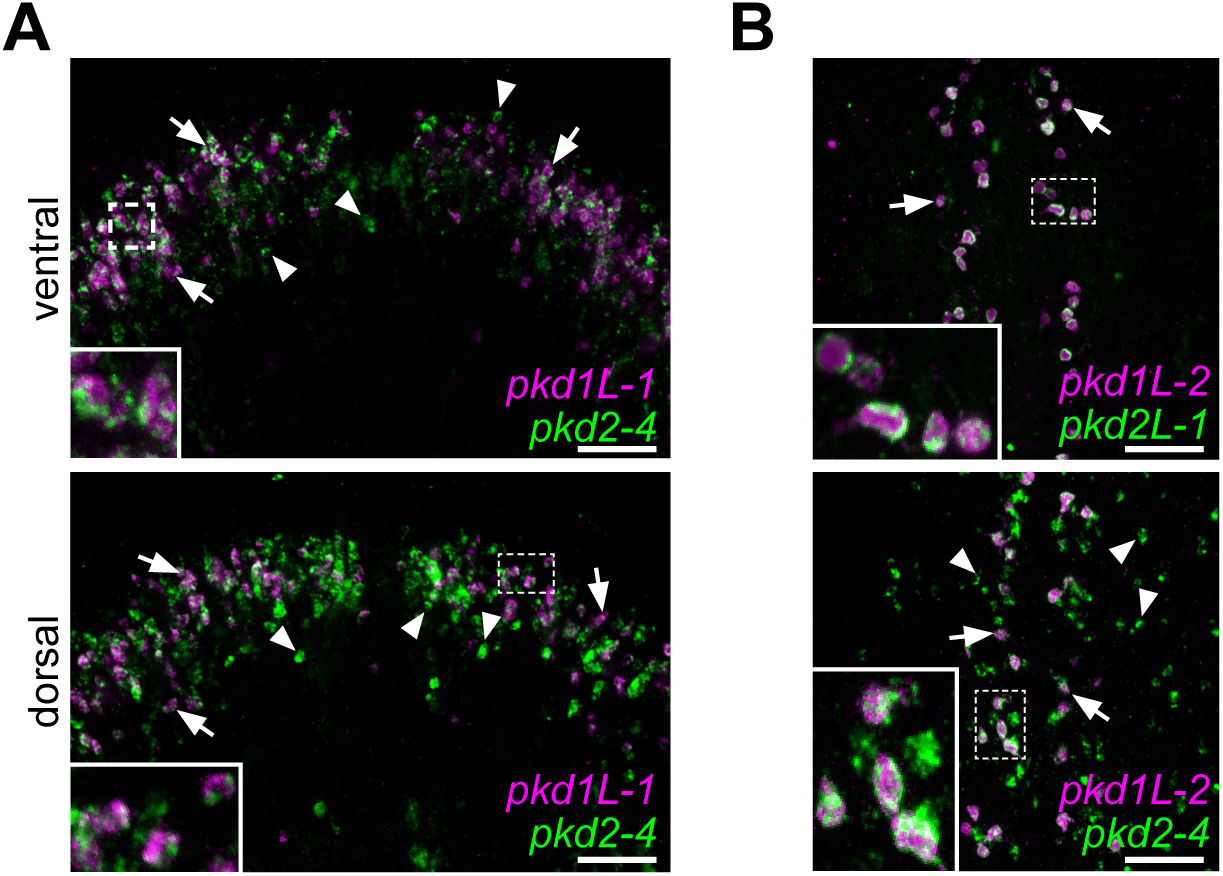
***pkd* genes that produced RNAi phenotypes are co-expressed in sensory neurons.** (A-B) dFISH on *pkd1* genes (magenta) and *pkd2* genes (green) that resulted in behavioral phenotypes. (A) dFISH co-labeling of the two *pkd* genes that resulted in significant reductions in chemosensory behaviors, *pkd1L-1* and *pkd2-4*, shows that the more discrete population of *pkd1L-1* cells are co-labeled with *pkd2-4* in the auricle and brain branch region in both the ventral and dorsal regions of the head periphery (examples highlighted with white arrows). There are many additional *pkd2-4^+^* cells that were not co-labeled with *pkd1L-1* (examples highlighted with white arrowheads). (B) dFISH of the *pkd1* gene that resulted in a mechanosensory phenotype, *pkd1L-2,* with both of the *pkd2* genes that resulted in mechanosensory phenotypes, *pkd2L-2* and *pkd2-4*, showed that *pkd1L-2^+^* cells are co-expressed with both *pkd2* mechanosensory genes (examples highlighted with white arrows). *pkd2-4^+^*cells represent a larger population that also includes *pkd1L-2-* cells (examples highlighted with white arrowheads). White dashed boxes indicate the higher magnification image insets. Scale bars, 50 µm, n ≥ 3 worms tested with all worms displaying similar expression patterns.

## Discussion

### *pkd* genes are expressed in neurons distributed across the body and pharynx of planarians

Previous work revealed that *pkd* genes are expressed in sensory regions and play roles in processes like sensing water flow or vibration in planarians. Here, we sought to characterize all *pkd* family members, including the *S. mediterranea* homologs, *pkd1L-1*, *pkd2-2*, and *pkd2-3,* that had yet to be analyzed. The planarian nervous system is surprisingly heterogeneous, with many specialized cell types that can be predominantly distinguished by unique gene expression signatures. Combined with previous reports, we conclude that *pkd* genes are exclusively expressed in neuronal cells in the asexual *S. mediterranea* biotype. This observation is consistent with previous reports indicating *pkd* genes are not expressed in the planarian ciliated excretory cell types, and we did not observe the types of phenotypes associated with protonephridia defects, such as osmoregulatory defects like bloating.

Because of the known roles of *pkd* genes in mating behaviors and reproductive processes in divergent organisms like *C. elegans, Drosophila*, sea urchins, and humans, curiosity drove us to examine the expression of *pkd* genes in the hermaphroditic strain of *S. mediterranea* (Fig. S3). The analysis yielded more questions than answers, opening the door to a future study to assess *pkd* genes’ specific cell-type expression and roles in mating behavior or fertilization. In addition to the neuronal patterns observed in asexual worms, we observed striking expression in reproductive organ anatomical regions of mature *S. mediterranea* hermaphrodites. However, we do not know the identities of the cells expressing *pkd* genes in the reproductive organs, which will require future experimentation to resolve if genes are expressed in sensory neurons or in reproductive somatic and gonadal cells. A recent study generated scRNA-seq data for the sexual biotype (Issigonis et al., 2024), so this new resource could advance the identification of cell types before performing double in situ hybridization experiments in hermaphrodites to provide insight into the cell-type expression of *pkd* genes. Interestingly, Pkd2 is found in sperm, and its function is required for fertilization (Gao et al., 2003; Watnick et al., 2003; Kierszenbaum, 2004). We observed *pkd2-3* expression in the planarian testes, suggesting it will be interesting to investigate whether it plays a conserved role in male fertility.

### *pkd* gene roles in sensory reception and behavior

We extended previous findings using behavioral assays showing that *pkd* genes have robust roles in mechanosensory and chemosensory reception (Ross et al., 2018). Our data supports that *pkd1L-2*, *pkd2-4*, and *pkd2L-1* are required for the mechanosensory function of the sensory population along the dorsal and peripheral ciliated stripes. Interestingly, gene co-expression analysis showed that these transcripts are expressed in the same neurons. Based on the canonical function of Polycystin-1 proteins, we surmise that *pkd1L-2* regulates the activity of *pkd2-4* and *pkd2L-*1 (Fedeles et al., 2014). We hypothesize that *pkd1L-2* acts as the receptor and couples the mechanical stimulation to either *pkd2-4* or *pkd2L-1* channels. Although double-FISH revealed the expression of both transcripts in double-positive cells, concluding that these proteins are co-localized or working together would require other approaches, like generating antibodies for protein immunostaining or biochemical assays.

Furthermore, we found that *pkd1L-1* and *pkd2-4* have roles in the chemosensory behaviors associated with feeding (Fig. 5). These genes were expressed in the brain branches and *pkd2-4* was also expressed in the auricle area, strongly indicating that cells within these structures are involved in chemosensation. Conversely, we did not observe phenotypes for *pkd2-2* and *pkd2-3*, which were specifically expressed in the pharynx. However, we did not directly test nor quantify pharyngeal behavior as in Miyamoto et al. (2020). Thus, it remains possible that pharyngeal *pkd* genes have roles in chemosensation. From our phenotype analysis, we can conclude that *pkd1L-1* and *pkd2-4* implicate head neurons in chemotaxis behaviors independent of the pharynx. *pkd1L-1* RNAi phenotypes were only observed in regenerates, suggesting stronger penetrance when the animals were challenged to fully regenerate heads.

We sought to redesign rigorous assays for straightforward set up and implementation (see Methods) that are based on our previous work (Ross et al., 2018); unfortunately, the code for the published assays was no longer available (the data was lost in a computer hardware failure; E.-M. S. Collins, personal communication). Our redesigned assays complement other existing assays to measure locomotion and other modalities like thermotaxis and thigmotaxis (Inoue and Agata, 2022). Because we only challenged the RNAi groups with assays related to mechanosensory and chemosensory abilities due to the defects seen in *soxB1-2* and downstream genes, other assays, such as thermosensation and thigmotaxis could reveal roles for some of the genes in this screen, especially the PKD2 family genes, which fall within the TRP family and have been demonstrated to function independently of PKD1 family members (Esarte Palomero et al., 2023). It has already been shown in *D. japonica* that *trpm3* participates in thermosensation (Inoue et al., 2014). It is possible *pkd* genes are involved in other sensory modalities like detection of chemical changes or temperature that have been shown to influence behaviors like reactions to water currents in other planarian species (Allen, 1915). In addition, further exploration using the scRNA-seq datasets for *S. mediterranea* could uncover other gene signatures that could help predict functions of the *pkd^+^* cell populations in the auricles or the pharynx.

### Concluding Remarks

This study sought to characterize the role of *pkd* genes in sensory neuron regeneration and function in the planarian *S. mediterranea*. In vertebrates, PKD1 and PKD2 genes are expressed in excretory organs and mutations in either protein causes Autosomal Dominant Polycystic Kidney Disease; however, studies have increasingly implicated PKD family genes in neural development and functions like taste reception (Harris and Torres, 2009; Ohata et al., 2015; England et al., 2017). In many invertebrates, including planarians, Pkd-like genes are expressed in nervous system cells (Barr et al., 2001; O’Hagan et al., 2014; McLaughlin, 2017; Fincher et al., 2018; Ross et al., 2018; Sebe-Pedros et al., 2018; Hulett et al., 2023; Sakagami et al., 2024). Consistent with those observations, *pkd* genes have strong expression in ciliated sensory neurons and have functions in sensory reception in *S. mediterranea*. Although additional evidence is required, it is tempting to speculate that an ancestral function of Pkd-like proteins is in sensory reception and is later co-opted in other organs like the vertebrate kidney. This work contributes to knowledge of *pkd* gene function in the Platyhelminthes, which will be useful for comparative studies on the evolution of neuronal functions.

## Acknowledgements

This study was supported by a California Institute for Regenerative Medicine Grant EDUC4-12813 postdoctoral fellowship to M.A. and NIH R01GM135657 to RMZ. We thank Dr. Yusuf Ozturk for help with the Arduino code; Lia Escober and Malia Huff for help collecting PKD homolog sequences for phylogenetic analysis. Portions of the assay design cartoons were created with BioRender.com.

## Methods

### Planarian care

Asexual planarians (CIW4) were maintained in 1X Montjuïc salts (1.6 mM NaCl, 1.0 mM CaCl_2_, 1.0 mM MgSO_4_, 0.1 mM MgCl_2_, 0.1 mM KCl, 1.2 mM NaHCO_3_) in plastic containers in an incubator at 20°C. Worms were fed homogenized liver and cleaned once per week. Planarians were starved one week prior to RNAi and fixation for in situ experiments.

### Cloning PKD genes

Nine PKD genes were previously identified by Thi-Kim Vu et al. (2015). We used the Conserved Domain Database to extract the PKD_Channel domain from *pkd1L-1* (dd_Smed_v6_13975_0_1) and *pkd2L-1* (dd_Smed_v6_ 15626 _0_1) (Wang et al., 2023) and then translated the domain sequences on Expasy (https://web.expasy.org/translate/). We entered these peptide sequences into the BLAST tool at PlanMine (Rozanski et al., 2019) and performed a TBLASTN search against the dd_Smed_v6 transcriptome with an expected e-value cut-off of e^-10^, which returned the nine PKD genes that were previously identified. The results were saved as Table S1. Genes fragments were cloned from cDNA or were purchased as pre-synthesized nucleotide eBlocks from Integrated DNA Technologies, https://www.idtdna.com/ (Table S2). Fragments were inserted into pPR242-T4P using ligation-independent cloning and transformed into HT115 (Adler and Alvarado, 2018). Primers, cDNA sequences, and eBlock fragment sequences for genes used in this paper are listed in Table S2.

### PKD protein domain comparison to human Polycystin proteins

*S. mediterranea pkd* transcript sequences and human PC*-*1 (P98161), PKD1L-1 (Q8TDX9), PKD1L-2 (Q7Z442), PKD1L-3 (Q7Z443), PC-2 (Q13563), PKD2L-1 (Q9P0L9), PKD2L-2 (Q9NZM6) protein sequences were uploaded into the legacy Pfam website, version 35 (pfam-legacy.xfam.org) using the search function with an expected-value cutoff set to 1.0. In all instances where transcripts were used, only one frame demonstrated significant hits to domains. The domain information was copied from Pfam, and representative images were created copying all domains represented in Pfam.

### Phylogenetic analysis

The software Geneious (www.geneious.com) was used to create a multiple alignment using the MUSCLE 5.1 plugin (Edgar, 2022). Protein alignments were manually inspected, and the PKD Channel Domains (aligned to the human Pfam PKD1 and PKD2 cation channel domains listed in Supplementary Files 1-2) were extracted for performing Bayesian inference of phylogeny using the MrBayes 3.2.6 (Huelsenbeck and Ronquist, 2001) plugin developed by Marc Suchard and the Geneious Team with the following settings: unconstrained branch length, shape parameter exponential of 10, 1.1Million chain length, 4 heated chains, 0.2 heated chain temp., WAG substitution model, gamma rate variation model, 10% burnin length with subsampling frequency of 200. All sequences used and the protein alignments are provided in Supplementary Files S1-4.

### Whole-mount in situ hybridization

Riboprobes were synthesized using either digoxigenin-11-UTP (DIG, Roche) or dinitrophenol-11-UTP (DNP, Perkin Elmer) as described in King, R. S. and Newmark (2013). Whole-mount in-situ hybridizations were performed on asexual planarians as previously described (King, R. S. and Newmark, 2013)39], except for the addition of a 1.5 or 3 minute incubation in a reduction solution (50 mM DTT, 1% NP-40, 0.5% SDS in 1X PBS) after fixation as in (Pearson et al., 2009). Hybridization and post-hybridization wash steps were performed at either 56°C or 58°C. Chromogenic asexual WISH samples were incubated with anti-DIG-AP (1:2000) and then developed with an NBT-BCIP solution (Roche) as previously described (King, R. S. and Newmark, 2013). Fixation and processing of sexual *S. mediterranea* for whole-mount in-situ hybridizations were performed as described in Issigonis et al. (2022) on worms approximately 8 mm in length that were checked visually for the presence of the gonopore prior to experimentation. The following exceptions were made to the protocol: animals were incubated in reduction solution (50 mM DTT, 1%NP-40, 0.5% SDS, in PBS) for 5 min at 37°C after being fixed. The hybridization protocol was performed as described in (King, R. S. and Newmark, 2013) except the hybridization time was increased to 36 hours, and four extra 0.1X SSCx (SSC + 0.1% Triton X-100) washes were performed post-hybridization. For WISH on isolated pharynxes, we followed a chemical amputation for pharynx removal as described in Miyamoto et al. (2020) and performed the fixation and WISH protocol as in King, R. S. and Newmark (2013). dFISH samples were fixed and hybridized in a mixture of the DIG and DNP probes, as described above. After SSCx washes, samples were washed twice in TNTx (0.1 M Tris pH 7.5, 0.15 M NaCl, 0.3% Triton X-100) at room temperature for 10 minutes, blocked in TNTx-blocking solution (5% heat-inactivated horse serum, 5% Roche Western Blocking Buffer in TNTx) and then incubated in anti-DNP-POD (Vector Laboratories, 1:2000) in TNTx-blocking solution overnight at 4°C. Samples were then washed six times for 20 minutes at room temperature in TNTx and then developed with Cy3-Tyramide as described in (King et al 2013). After MABTw (Maleic Acid Buffer + 0.1% Tween-20) washes, animals were incubated in MABTw blocking solution (5% heat-inactivated horse serum, 5% Roche Western Blocking Buffer in MABTw) for 2 hours at room temperature and then incubated in anti-DIG-AP (Roche, 1:2000) overnight at 4°C and washed and developed with Fast Blue as described in (King 2013). After post-development washes, samples were quenched with CuSO4 solution (10mM CuSO4, 50mM ammonium acetate pH5.0) for 1 hour at room temperature, washed in ultrapure water two times for 5 minutes at room temperature and then with PBSTx 2 times for 5 minutes at room temperature, and incubated overnight in 5 ug/ml DAPI in PBSTx at 4°C. Worms were mounted in 80% glycerol or a 1:1 mixture of 80% glycerol and Vectashield under a No. 1 thickness cover glass and imaged on a Leica M250C stereomicroscope fitted with a Leica DFC 450 color camera for chromogenic WISH or Leica Stellaris 5 Confocal microscope running LAS X v4.6.1 with a 20x/0.75 dry objective. dFISH images were acquired through the Z-plane of the region of interest at the optimal interval identified by the software and further processed to extract slices of interest, create a maximum intensity projection, and overlay multiple pseudocolored channels in either FIJI (ImageJ2 version 2.9.0) (Schindelin et al., 2012) or LAS X.

### RNA interference

Double-stranded RNA was synthesized by bacterial expression (Adler and Alvarado, 2018) or *in vitro* transcription reactions using the MEGAscript T7 RNA synthesis kit (Invitrogen, Inc) as described in (Rouhana et al., 2013). For *in vitro* dsRNA, the synthesized dsRNA was diluted to a concentration of 1 µg/µl and stored in DEPC-treated water at −20°C. The pellets were freshly prepared by mixing the liver extracts with an agarose solution (1.25 µl of Liver extract + 0.25 µl of 2% low-melting agarose and 0.5 µl of dsRNA per planarian in the dish) and then frozen for approximately one hour before feeding. For the non-extended feeding protocol, planarians were fed either bacterially expressed or *in vitro* transcribed dsRNA liver pellets eight times over 4 weeks. Extended-feed RNAi planarians were fed 14 times over 7 weeks.

### Vibration assay

Vibration assays were performed the day following completion of RNAi feeding as well as on planarians that regenerated their heads 13-15 days following amputation anterior to the pharynx. Groups of 5 planarians were placed near the center of a 100 mm diameter petri dish filled with 40 mL of Montjuïc salts that was anchored with silicone paste into a plate lid fixed onto a cold LED board. The setup is illustrated in (Fig. S4C-C’). After worms were gliding normally, a tapping program (code in Supplementary File S5) running from a microcontroller board (Arduino Uno Rev3) connected to a Seamuing MG996R Micro Servo motor (Model No. 5123164-2360-1341090661) with an attached plastic arm that was 112 mm in length from the center of the rotor to the end of the arm, 4 mm thick, and 8 mm wide (obtained from Fielect 75 Type Plastic Gears Kit FLT20191223S-1008 from amazon.com). The end of the arm, wherein the tapper struck the dish, was wrapped with laboratory tape to increase the thickness to approximately 8 mm in thickness. Taping also served to dampen the vibration of the arm during the tapping stimulus. The mechanism was programmed to deliver five taps at a rate of one tap every 75 ms to the side of the dish. Supplementary File S5 provides a link to the code used to control a Servo motor from an Arduino board (Supplementary File S5).

### Rheosensation assay

Rheosensory assays (Fig. S4E) were performed the day following completion of RNAi feeding and on planarians that regenerated their heads 13-15 days following amputation anterior to the pharynx. A single asexual worm was placed inside a clear plastic chamber with dimensions of 90 mm in length by 40 mm in height filled with 10 ml of 1X Montjuïc salts and observed until gliding normally and parallel to the front of the chamber, in some cases, the planarian would need to be pipetted with a transfer pipet and moved to help orient them back into the middle of the chamber. Then, 30 μl of ultrapure water containing yellow food dye was ejected on the dorsal side of the worm using a Finnpipette P50 (50 μl) pipette.

### Chemosensation assay

Food pellets were prepared by making a gelled liver paste containing a ratio of 25 µl of liver paste (500 µl of pureed liver mixed with 460 µl of ultrapure water and 40 µl red food coloring) to 5 µl of 2% low melting point agarose and then pipetting 15 µl drops of gelled liver paste onto wax paper which were then frozen at −20°C for at least 1 hr. Chemosensory assays were performed in a clear plastic STORi 6” x 3” X 2” stackable bin (purchased from amazon.com). The dimensions of the bottom of the rectangular tray were 140 mm x 65 mm and contained marks for a ‘start zone’ line 10 mm away from the end of the tray and a food line 80 mm away from the ‘start zone’ line with lines 5 mm from either side of the food line to denote the ‘food zone’ (see Figure S5 for tray illustration). The tray was filled with 50 ml of 1X Montjuïc salts, and then three of the gelled liver paste pellets were placed across the food zone line. The food was allowed three minutes of diffusion time in the water, and then 10 worms from either the control or experimental group were transferred into the ‘start zone’ and allowed a 10-minute time frame to glide 2/3 distance across a 140 mm dish to a 10 mm ‘food zone’ where 3 agarose-liver pellets were placed, and where chemical signals emanating from the food pellets would be strongest (Fig. S5A). We analyzed videos of the worms during the 10-minute assay and measured both the total time that planarians spent within the ‘food zone’ as well as the total number of worms that consumed food during that period for both intact and regenerate worms (RNAi scheme summarized in Fig. S4B). Worms were visually inspected for red food dye, and the number of worms that ate were recorded in addition to the analysis described below.

### Behavioral assay digital recording and video analysis

All behavioral assays were recorded using a Basler Ace 2 Pro ac1440-220uc camera recording onto a PC running Basler’s pylon Viewer 64-bit version 6.3.0 software at a frame rate of 10 frames/second and frame size of 1440 x 1080 pixels. *gfp*(*RNAi*) planarians were used as the control group for all experiments. Video frames were analyzed in FIJI (ImageJ2 version 2.9.0) (Schindelin et al., 2012). For Vibration and Rheosensory assays, the line tool in FIJI was used to measure the longest pre-stimulus length of each gliding worm and the length of the worm following application of the stimulus. The percent change in the length was calculated as Percent Change = [(Length_Pre-Stimulus_ – Length_Post-Stimulus_) / Length_Pre-Stimulus_] X 100. For the chemosensory assay, the worms were tracked visually one at a time within frames to determine every timepoint when that worm entered and exited the food zone. The sum of all time (in seconds) spent in the food zone during the 10-minute assay period was calculated. For all behavioral experiments, measurements were collected for at least 10 worms per experimental group over at least two independent experiments.

### Statistical analysis

All statistical analyses were performed in GraphPad Prism 9, and graphs were created in the same software. One-way ANOVA was performed and corrected for multiple comparisons using Dunnett’s correction. All means were compared to a *gfp* control group. Statistical significance was accepted at values of p <0.01.

### Supplementary Material

**Figure S1.**
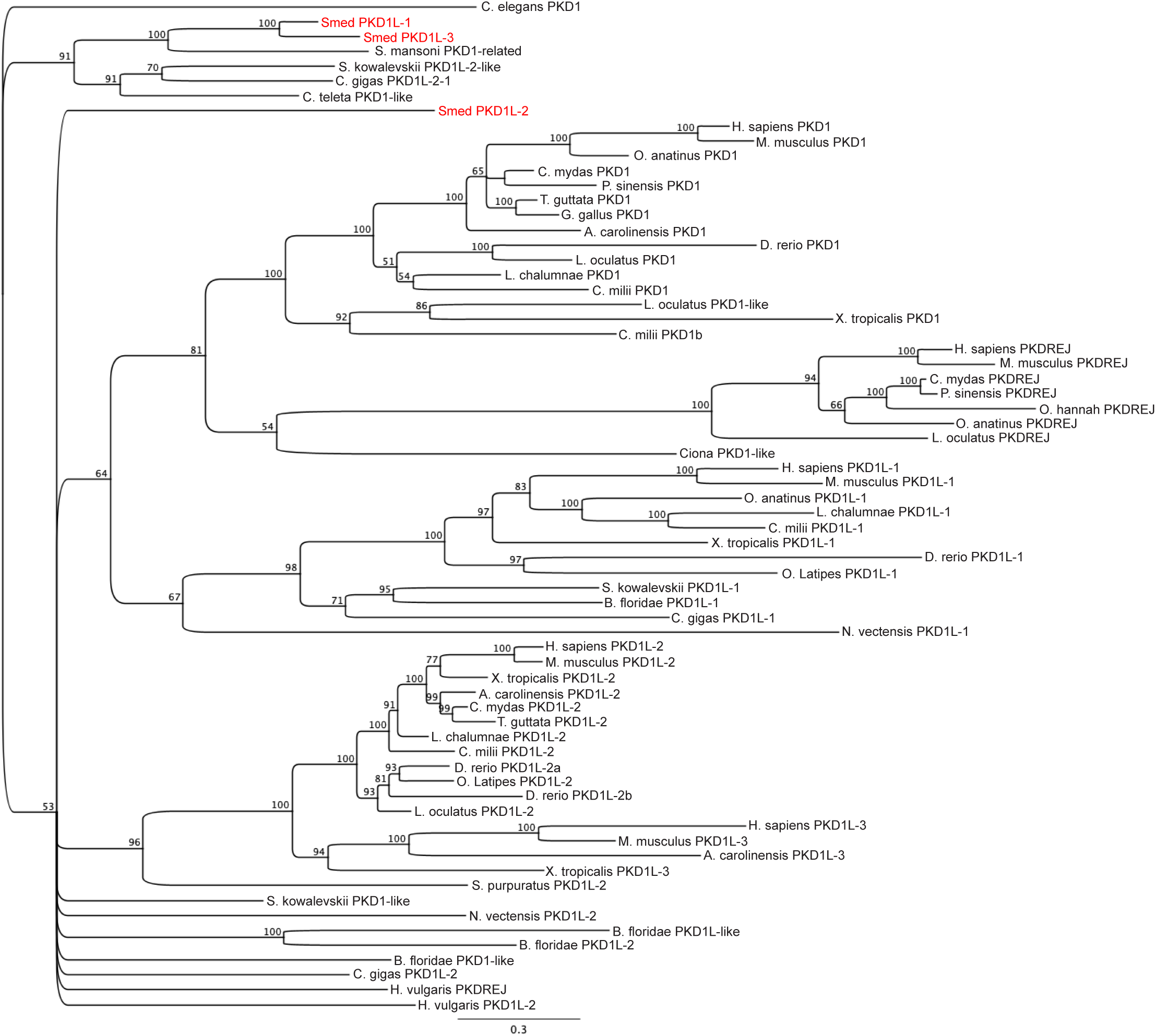
A Bayesian inference phylogeny of PKD1 and PKD1L proteins. Node support values are listed as percent support next to the relevant node.

**Figure S2.**
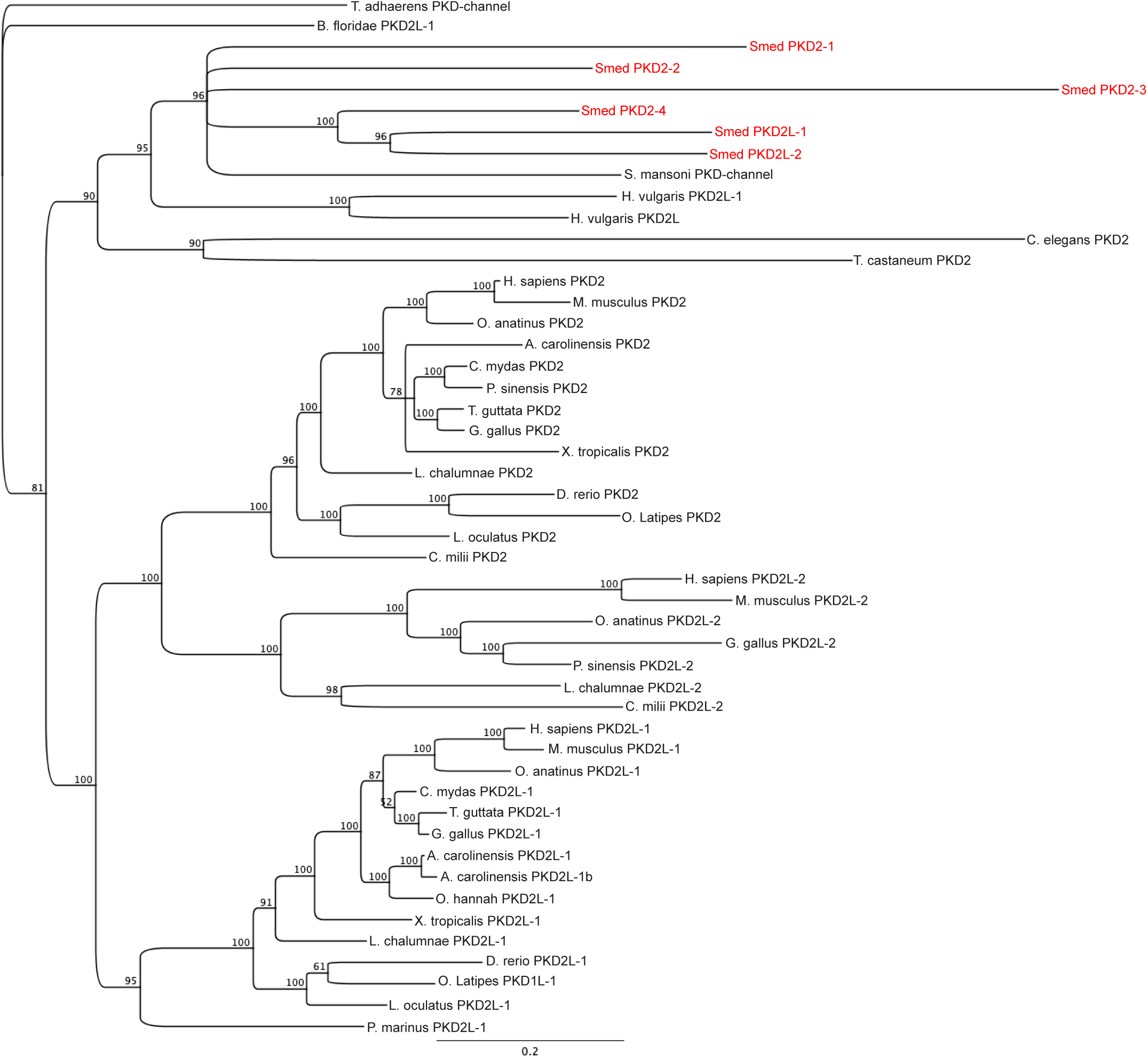
A Bayesian inference phylogeny of PKD2 proteins. Node support values are listed as percent support next to the relevant node.

**Figure S3.**
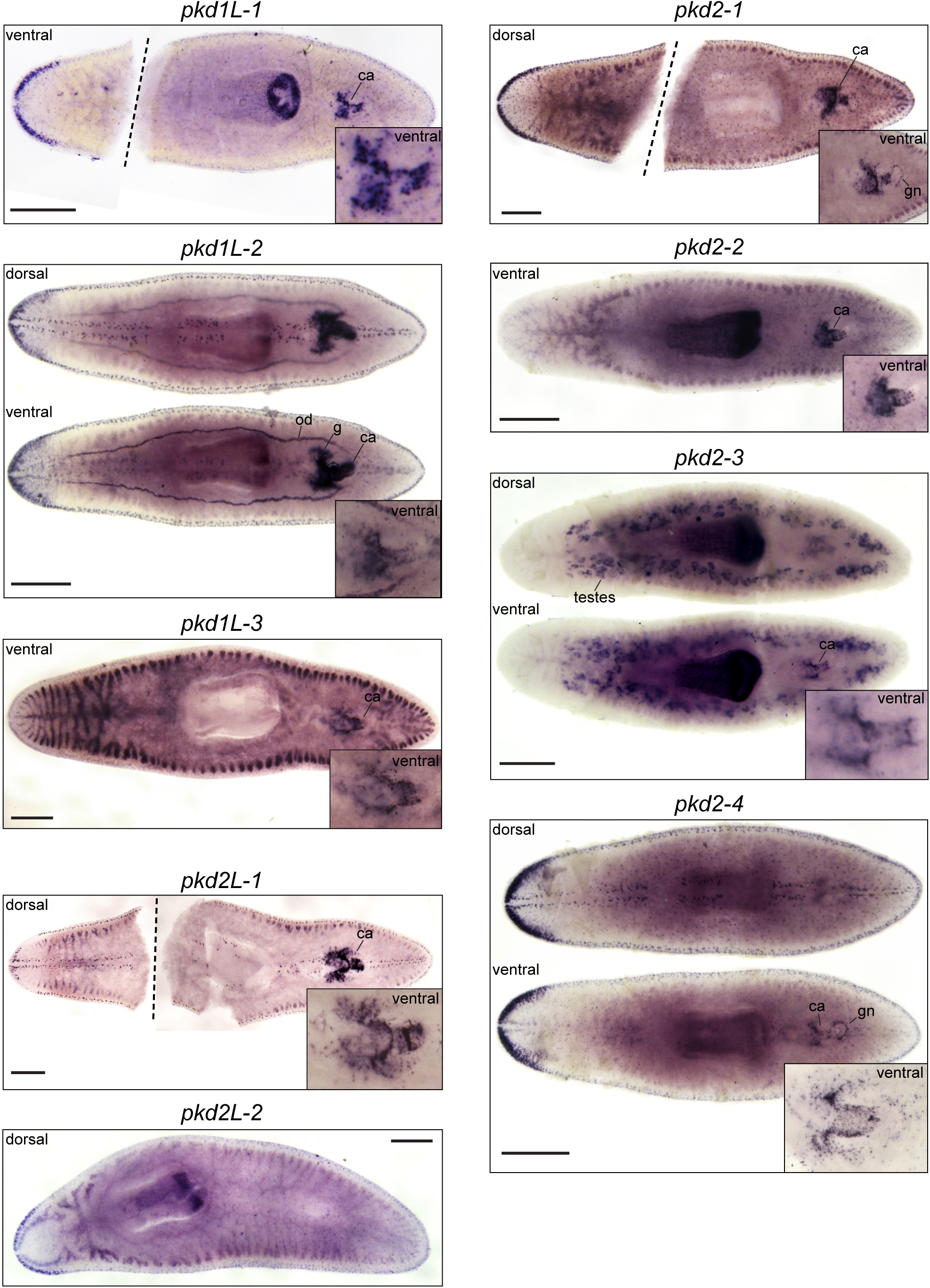
In situ hybridization to *pkd* genes in *S. mediterranea* hermaphrodites. The expression of *pkd* genes mirrors in situ patterns in the asexual strain. However, eight out of nine planarian *pkd* genes (all except *pkd2L-*2 in the bottom left panel) showed striking expression associated with reproductive structures: copulatory apparatus, ca; oviduct, od; gland cells, g; testes; and the gonopore, gn. Inset showed higher magnification of the copulatory apparatus regions for all genes detected. Scale bars = 500 µm.

**Figure S4.**
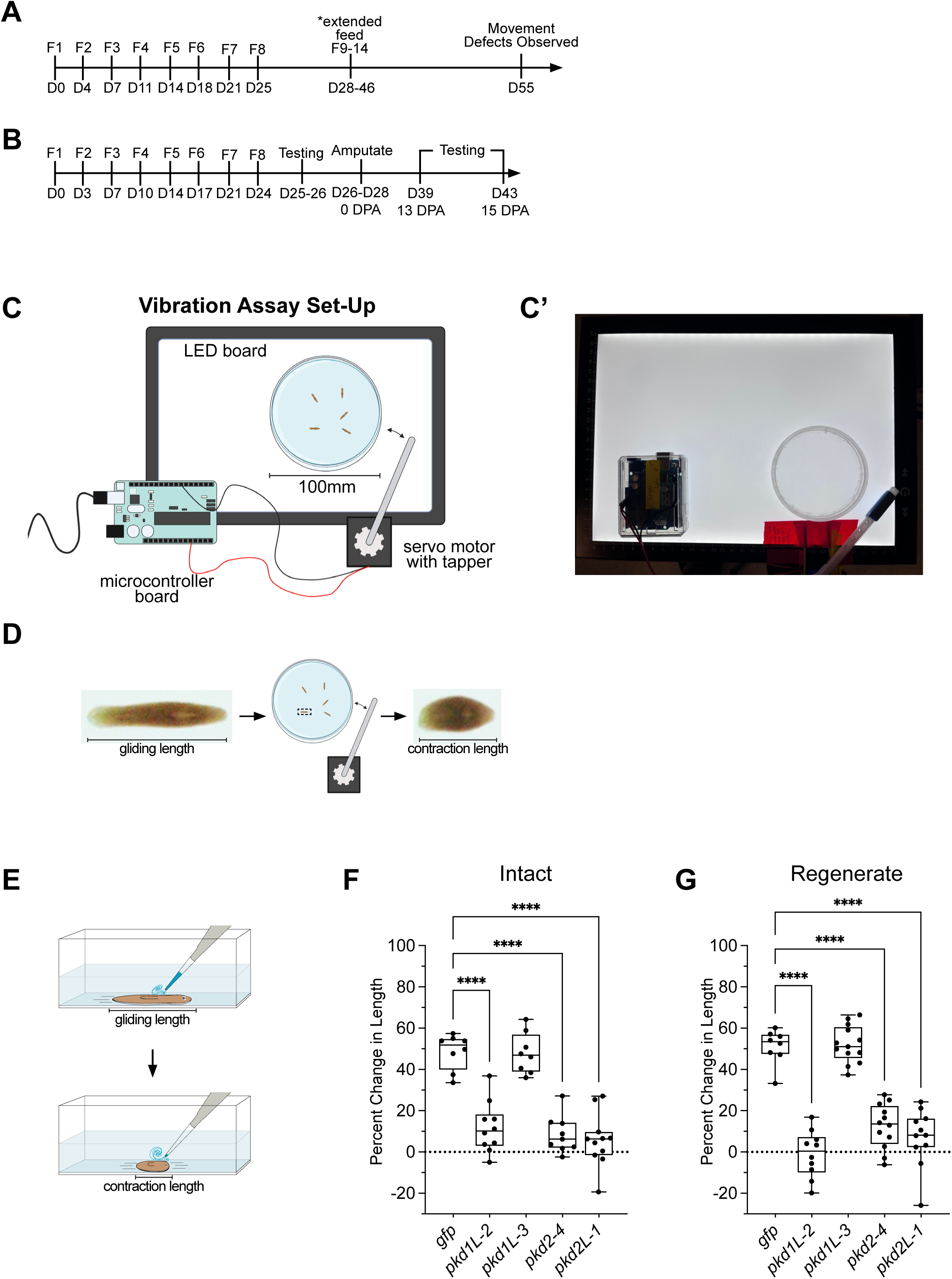
Timelines for RNAi experiments and vibration assay. (A) Timeline for *pkd* RNAi experiments and subsequent behavioral testing. All testing was performed between 13- and 15-days following amputation. F, feed; D, day in an experiment; DPA, days post amputation. (C-E) depictions of the mechanosensory assays. (C) An illustration of the tapping device used to test mechanosensation is in Figure 4 (C’). A photo of the tapping device setup is shown. (D-E) Illustrations and still images from behavioral videos showing how pre-stimulus “gliding” length and post-stimulus “contraction” length were measured for the rheosensory (D) and vibration (E) assays. The worms depicted are examples of a typical wild-type behavioral response. (F-G) Worms with *pkd* knockdowns that demonstrated a significant loss of mechanosensory function in the vibration assay were also tested using the rheosensory assay to validate the observed loss of mechanosensation in intact (F) and regenerate (G) worms. The percent change in length was calculated and plotted in a box plot wherein the box extends from the 25^th^ to 75^th^ percentiles of the range, and the bisecting line represents the median. The whiskers extend the full range of the data. *pkd1L-3*, which did not display a mechanosensory phenotype, was included as an additional negative control. N = 8-13 worms for each experimental group. ^∗∗∗∗^p < 0.0001; all other groups are non-significant (p > 0.01); One-way ANOVA with Dunnett’s multiple comparisons test.

**Figure S5.**
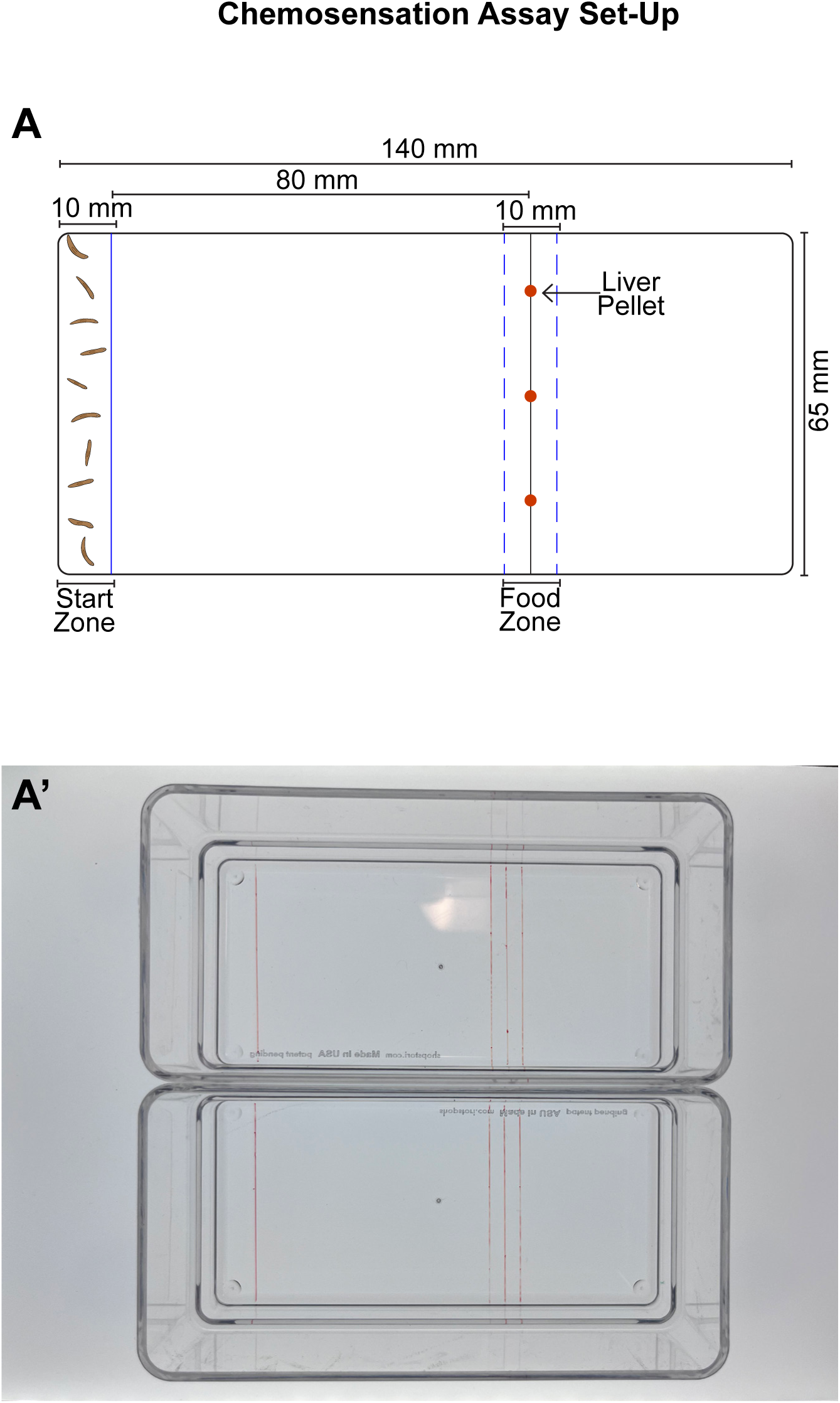
Chemosensation assay. (A) Illustration of the custom-made chamber used for chemosensation assays displaying the markings on the dish used to determine inclusion in the ‘feeding zone’ and start zone as explained in the methods. The lines were drawn on the bottom of the clear dish and could be visualized in the videos, which were taken with an overhead camera. (A’) Photo of trays with drawn lines used in this study.

**Table S1.** Results from BLAST of PKD domain to Planarian Transcriptome.

**Table S2.** Primers and eBlock sequences used in this study.

**File S1**. Protein sequences used for PKD1 phylogenetic analysis in Figure S1.

**File S2.** Protein sequences used for PKD2 phylogenetic analysis in Figure S1.

**File S3.** PKD1 cation channel alignment file used for phylogenetic analysis in Figure S1.

**File S4.** PKD2 cation channel alignment file used for phylogenetic analysis in Figure S2.

**File S5.** Link to Arduino code for controlling tapping device with servo motor

**Video S1.** Control worms 28 days following the first RNAi feeding display normal gliding locomotion movements in the arena used for the chemosensory assay.

**Video S2.** *pkd2-4(RNAi)* worms 28 days following the first RNAi feeding display normal gliding locomotion movements in the arena used for the chemosensory assay.

**Video S3.** Control worms at 55 days following the first RNAi feeding display normal gliding locomotion movements.

**Video S4.** *pkd2-4(RNAi)* worms 55 days following the first RNAi feeding display slow, inching, jerky locomotion movements.

## References

Adler, C.E., and Alvarado, A.S. (2018). Systemic RNA Interference in Planarians by Feeding of dsRNA Containing Bacteria. Methods Mol Biol 1774, 445–454.

Allen, G.D. (1915). REVERSIBILITY OF THE REACTIONS OF PLANARIA DOROTOCEPHALA TO A CURRENT OF WATER. The Biological Bulletin 29, 111-[128]-111.

Almazan, E.M.P., Ryan, J.F., and Rouhana, L. (2021). Regeneration of Planarian Auricles and Reestablishment of Chemotactic Ability. Front Cell Dev Biol 9, 777951.

Arnold, C.P., Lozano, A.M., Mann, F.G., Jr., et al. (2021). Hox genes regulate asexual reproductive behavior and tissue segmentation in adult animals. Nat Commun 12, 6706.

Barr, M.M., DeModena, J., Braun, D., et al. (2001). The Caenorhabditis elegans autosomal dominant polycystic kidney disease gene homologs lov-1 and pkd-2 act in the same pathway. Curr Biol 11, 1341–1346.

Edgar, R.C. (2022). Muscle5: High-accuracy alignment ensembles enable unbiased assessments of sequence homology and phylogeny. Nat Commun 13, 6968.

England, S.J., Campbell, P.C., Banerjee, S., et al. (2017). Identification and Expression Analysis of the Complete Family of Zebrafish pkd Genes. Front Cell Dev Biol 5, 5.

Esarte Palomero, O., Larmore, M., and DeCaen, P.G. (2023). Polycystin Channel Complexes. Annu Rev Physiol 85, 425–448.

Farnesi, R.M., and Tei, S. (1980). Dugesia lugubris s.l. auricles: research into the ultrastructure and on the functional efficiency. Riv Biol 73, 65–77.

Fedeles, S.V., Gallagher, A.R., and Somlo, S. (2014). Polycystin-1: a master regulator of intersecting cystic pathways. Trends Mol Med 20, 251–260.

Fincher, C.T., Wurtzel, O., de Hoog, T., et al. (2018). Cell type transcriptome atlas for the planarian Schmidtea mediterranea. Science 360.

Fraenkel, A., Gunn, D. L. (1961). The orientation of animals. New York: Dover.

Gao, Z., Ruden, D.M., and Lu, X. (2003). PKD2 cation channel is required for directional sperm movement and male fertility. Curr Biol 13, 2175–2178.

Grohme, M.A., Schloissnig, S., Rozanski, A., et al. (2018). The genome of Schmidtea mediterranea and the evolution of core cellular mechanisms. Nature 554, 56–61.

Harris, P.C., and Torres, V.E. (2009). Polycystic kidney disease. Annu Rev Med 60, 321–337.

Horio, N., Yoshida, R., Yasumatsu, K., et al. (2011). Sour taste responses in mice lacking PKD channels. PLoS One 6, e20007.

Huelsenbeck, J.P., and Ronquist, F. (2001). MRBAYES: Bayesian inference of phylogenetic trees. Bioinformatics 17, 754–755.

Hulett, R.E., Gehrke, A.R., Gompers, A., et al. (2023). A wound-induced differentiation trajectory for neurons. bioRxiv.

Hulett, R.E., Rivera-Lopez, C., Gehrke, A.R., et al. (2024). A wound-induced differentiation trajectory for neurons. Proc Natl Acad Sci U S A 121, e2322864121.

Inoue, T., and Agata, K. (2022). Quantification of planarian behaviors. Dev Growth Differ 64, 16–37.

Inoue, T., Yamashita, T., and Agata, K. (2014). Thermosensory signaling by TRPM is processed by brain serotonergic neurons to produce planarian thermotaxis. J Neurosci 34, 15701–15714.

Ishimaru, Y., Inada, H., Kubota, M., et al. (2006). Transient receptor potential family members PKD1L3 and PKD2L1 form a candidate sour taste receptor. Proc Natl Acad Sci U S A 103, 12569–12574.

Issigonis, M., Browder, K.L., Chen, R., et al. (2024). A niche-derived nonribosomal peptide triggers planarian sexual development. Proc Natl Acad Sci U S A 121, e2321349121.

Issigonis, M., and Newmark, P.A. (2019). From worm to germ: Germ cell development and regeneration in planarians. Curr Top Dev Biol 135, 127–153.

Issigonis, M., Redkar, A.B., Rozario, T., et al. (2022). A Kruppel-like factor is required for development and regeneration of germline and yolk cells from somatic stem cells in planarians. PLoS Biol 20, e3001472.

Ivankovic, M., Haneckova, R., Thommen, A., et al. (2019). Model systems for regeneration: planarians. Development 146.

Kierszenbaum, A.L. (2004). Polycystins: what polycystic kidney disease tells us about sperm. Mol Reprod Dev 67, 385–388.

King, H.O., Owusu-Boaitey, K.E., Fincher, C.T., et al. (2024). A transcription factor atlas of stem cell fate in planarians. Cell Rep 43, 113843.

King, R.S., and Newmark, P.A. (2013). In situ hybridization protocol for enhanced detection of gene expression in the planarian Schmidtea mediterranea. BMC Dev Biol 13, 8.

Koehler, O. (1932). Beiträge zur Sinnesphysiologie der Süßwasserplanarien. Z. vergl. Physiol. 16, 606–756.

Liu, C., and Montell, C. (2015). Forcing open TRP channels: Mechanical gating as a unifying activation mechanism. Biochem Biophys Res Commun 460, 22–25.

LopezJimenez, N.D., Cavenagh, M.M., Sainz, E., et al. (2006). Two members of the TRPP family of ion channels, Pkd1l3 and Pkd2l1, are co-expressed in a subset of taste receptor cells. J Neurochem 98, 68–77.

Maser, R.L., Calvet, J.P., and Parnell, S.C. (2022). The GPCR properties of polycystin-1-A new paradigm. Front Mol Biosci 9, 1035507.

McLaughlin, S. (2017). Evidence that polycystins are involved in Hydra cnidocyte discharge. Invert Neurosci 17:1.

Miyamoto, M., Hattori, M., Hosoda, K., et al. (2020). The pharyngeal nervous system orchestrates feeding behavior in planarians. Sci Adv 6, eaaz0882.

Morris, Z., Sinha, D., Poddar, A., et al. (2019). Fission yeast TRP channel Pkd2p localizes to the cleavage furrow and regulates cell separation during cytokinesis. Mol Biol Cell 30, 1791–1804.

O’Hagan, R., Wang, J., and Barr, M.M. (2014). Mating behavior, male sensory cilia, and polycystins in Caenorhabditis elegans. Semin Cell Dev Biol 33, 25–33.

Ohata, S., Herranz-Perez, V., Nakatani, J., et al. (2015). Mechanosensory Genes Pkd1 and Pkd2 Contribute to the Planar Polarization of Brain Ventricular Epithelium. J Neurosci 35, 11153–11168.

Pearson, B.J., Eisenhoffer, G.T., Gurley, K.A., et al. (2009). Formaldehyde-based whole-mount in situ hybridization method for planarians. Dev Dyn 238, 443–450.

Plass, M., Solana, J., Wolf, F.A., et al. (2018). Cell type atlas and lineage tree of a whole complex animal by single-cell transcriptomics. Science 360.

Reddien, P.W. (2018). The Cellular and Molecular Basis for Planarian Regeneration. Cell 175, 327–345.

Roberts-Galbraith, R.H., Brubacher, J.L., and Newmark, P.A. (2016). A functional genomics screen in planarians reveals regulators of whole-brain regeneration. Elife 5.

Ross, K.G., Currie, K.W., Pearson, B.J., et al. (2017). Nervous system development and regeneration in freshwater planarians. Wiley Interdiscip Rev Dev Biol 6.

Ross, K.G., Molinaro, A.M., Romero, C., et al. (2018). SoxB1 Activity Regulates Sensory Neuron Regeneration, Maintenance, and Function in Planarians. Dev Cell 47, 331–347 e335.

Rouhana, L., Weiss, J.A., Forsthoefel, D.J., et al. (2013). RNA interference by feeding in vitro-synthesized double-stranded RNA to planarians: methodology and dynamics. Dev Dyn 242, 718–730.

Rozanski, A., Moon, H., Brandl, H., et al. (2019). PlanMine 3.0-improvements to a mineable resource of flatworm biology and biodiversity. Nucleic Acids Res 47, D812–D820.

Sakagami, T., Watanabe, K., Hamada, M., et al. (2024). Structure of putative epidermal sensory receptors in an acoel flatworm, Praesagittifera naikaiensis. Cell Tissue Res 395, 299–311.

Schindelin, J., Arganda-Carreras, I., Frise, E., et al. (2012). Fiji: an open-source platform for biological-image analysis. Nat Methods 9, 676–682.

Sebe-Pedros, A., Saudemont, B., Chomsky, E., et al. (2018). Cnidarian Cell Type Diversity and Regulation Revealed by Whole-Organism Single-Cell RNA-Seq. Cell 173, 1520–1534 e1520.

Thi-Kim Vu, H., Rink, J.C., McKinney, S.A., et al. (2015). Stem cells and fluid flow drive cyst formation in an invertebrate excretory organ. Elife 4.

Van Goethem, E., Silva, E.A., Xiao, H., et al. (2012). The Drosophila TRPP cation channel, PKD2 and Dmel/Ced-12 act in genetically distinct pathways during apoptotic cell clearance. PLoS One 7, e31488.

Wang, J., Chitsaz, F., Derbyshire, M.K., et al. (2023). The conserved domain database in 2023. Nucleic Acids Res 51, D384–D388.

Watnick, T.J., Jin, Y., Matunis, E., et al. (2003). A flagellar polycystin-2 homolog required for male fertility in Drosophila. Curr Biol 13, 2179–2184.

Wurtzel, O., Cote, L.E., Poirier, A., et al. (2015). A Generic and Cell-Type-Specific Wound Response Precedes Regeneration in Planarians. Dev Cell 35, 632–645.

