## Supplementary File S5 for "The role of *polycystic kidney disease-like* homologs in planarian nervous system regeneration and function": File S5. Arduino_tap_code_final.ino

```
#include <Servo.h>
#include <avr/wdt.h>


Servo myservo;  
#define servoPin 9 

int numberOfTaps = 5;
int lowerAngle   = 0; 
int neutralAngle = -30;
int higherAngle =  15; 
int delayBetweenTaps = 7000 ; 
int tapSpeed = 75; // time between taps , msec

int pos = 0; //tells servo to reset angle to 0 
int i = 0;


void setup() {
  myservo.attach(servoPin);
  wdt_enable(WDTO_8S);
  //myservo.writeMicroseconds(1000);  // set servo to mid-point


}


void loop() {
 
i = 0; 
while( i < numberOfTaps) { //tells the servo to do 5 total taps
 
    wdt_reset();          
    delay(tapSpeed); //delays for 75 ms
    myservo.write(higherAngle);    
    delay(tapSpeed);
    myservo.write(neutralAngle);             
 
  i = i+1; //repeats the process

  }
 
  i = 0;
  delay(tapSpeed);
  myservo.write(neutralAngle);              

  delay(delayBetweenTaps); //waits 15 seconds before repeating loop
}
```
